# Combined MD and QM/MM Calculations reveal Allostery Driven Promiscuity in Dipeptide Epimerases of Enolase Family

**DOI:** 10.1101/2022.05.01.490185

**Authors:** Ankita Tripathi, Kshatresh Dutta Dubey

## Abstract

The adaptability of the active site to amplify the secondary function is supposed to be the fundamental cause of the promiscuity and the evolution of new functions in the enzymes. In most cases, mutations occur close to the active site and/or in the catalytic site to change the active site plasticity to accommodate the non-native substrate. In the present study, using MD simulations and hybrid QM/MM calculations, we have shown a new way to enhance the promiscuity, i.e., the allostery-driven promiscuity. Using a case study of the AEE enzyme where the capping loop recognizes the substrate, herein, we show that a single site mutation (D321G) far from the capping loop can induce a large conformational change in the capping loop to recognize different substrates for different functions. The QM/MM calculations for the WT and mutated enzyme provide a first validation of the mechanism of 1,1-proton transfer and dehydration by the AEE enzyme. Since AEE epimerase possesses a highly conserved TIM-barrel fold, we believe that our study provides a crucial lead to understanding the mechanism of emergence of secondary function which can be useful to repurpose ancient enzymes for modern usage.

## 1. Introduction

Enzymes are nature’s ultimate machinery to catalyze some complex and unorthodox reactions.^1,2^ These enzymes are very efficient, but at the same time, they have evolved to catalyze a specific function which creates an initial roadblock to harnessing the catalytic efficiency of the natural enzymes for modern purposes.^3,4^ Therefore, tweaking enzymes for desired functions has become a holy grail for biochemists.^5^ However, the recent developments in directed evolutions and site-directed mutagenesis provide ample examples that bioengineering of some ‘strategic’ residues can evolve a specific enzyme for promiscuous functions.^6–9^ Therefore, the mechanistic study for such evolution of new (or secondary functions) other than the native function of a natural enzyme can provide a positive lead to repurpose ancient enzymes for the modern functions using bioengineering tools.^10–13^

So far, it is conventional to believe that the adaptive plasticity of the active site to accommodate secondary (or new) substrate may lead to the mutations and thus causes the evolution of promiscuous functions.^14–18^ However, there are a few examples where the mutation is far from the site of the conformational plasticity, and such conformational changes may allosterically trigger the promiscuity in the targeted enzyme.^19–21^ Dipeptide epimerase, a member of the enolase superfamily, offers such an example where a single site mutation in a *non-catalytic* residue of a specialist enzyme causes a spontaneous emergence of secondary activity shown by another specialist enzyme.^22^ For instance, Ala-Glu-Epimerase (AEE), the first functionally characterized epimerase, *specifically* catalyzes the 1,1-proton transfer reaction. Likewise, O-succinyl benzoate synthases (OSBS), another member of the enalose family, *specifically* catalyze dehydration reactions.^23–25^ Interestingly, both enzymes show just 23% overall sequence similarity.^26^ As can be seen from Figure 1, the active site residues in both enzymes remain very similar, however, a Glycine residue of OSBS enzyme close to the active site is replaced by an Aspartic acid in AEE (c.f. the position of G288 and D321 in Figure 1). Therefore, it may be perceived that Nature has replaced an Aspartate residue in AEE to evolve it for 1-1-proton transfer reactions during the evolution from a common progenitor of AEE and OSBS enzyme. This was further supported by a site-directed mutation study by Schmidt and co-workers where they have shown that a single mutation of D297→G (in *E. coli* where D297 is an analogue of D321 of *Bacillus Subtilis*), in the AEE enzyme causes a significant promiscuous activity for the OSBS enzyme at the cost of its native activity, however, a precise mechanism of the origin of promiscuity and loss of native activity in AEE is less elucidated till date.^22,26^ In a separate study by Schmidt et al, the role of capping loop located at the tip of substrate entrance channel, has been emphasized during the substrate recognition by comparing the Apo and Holo enzyme structures.^26^ However, due to the lack of experimental structure of the D297G mutant, the mechanism of the origin of the promiscuity is yet not clear. Furthermore, the question whether capping loop also plays a role in promiscuity due to the D297G mutation (D321G in *Bacillus Subtilis*) is not elucidated and will be intriguing to study.

**Figure 1.**
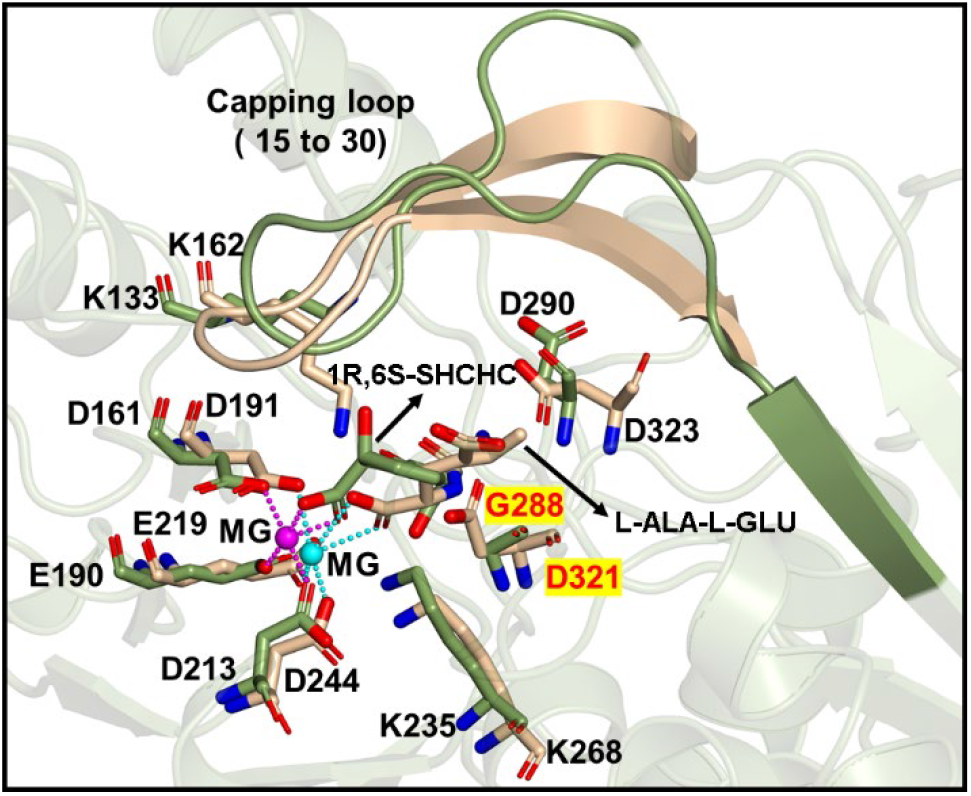
Superimposed crystal structures of AEE (PDB id: 1TKK) and OSBS (PDB id: 1R6W) enzymes. The active site residues are shown in stick presentations. AEE residues are shown in golden color while OSBS residues are in green color.

The previously *proposed* reaction mechanism for the 1-1 proton transfer by the AEE enzyme and dehydration by OSBS further entangles the mechanism of promiscuity.^26,27^ As can be seen from Scheme 1a and 1b, both enzymes catalyze the reaction through an enediolate anion intermediate where an active site base abstracts the α-proton from the substrate. However, the question that how the mutation of Asp321 to Glycine which itself doesn’t participate in any of the reactions causes an OSBS activity in AEE remains unclear from the above mechanistic scheme.

**Scheme 1.**
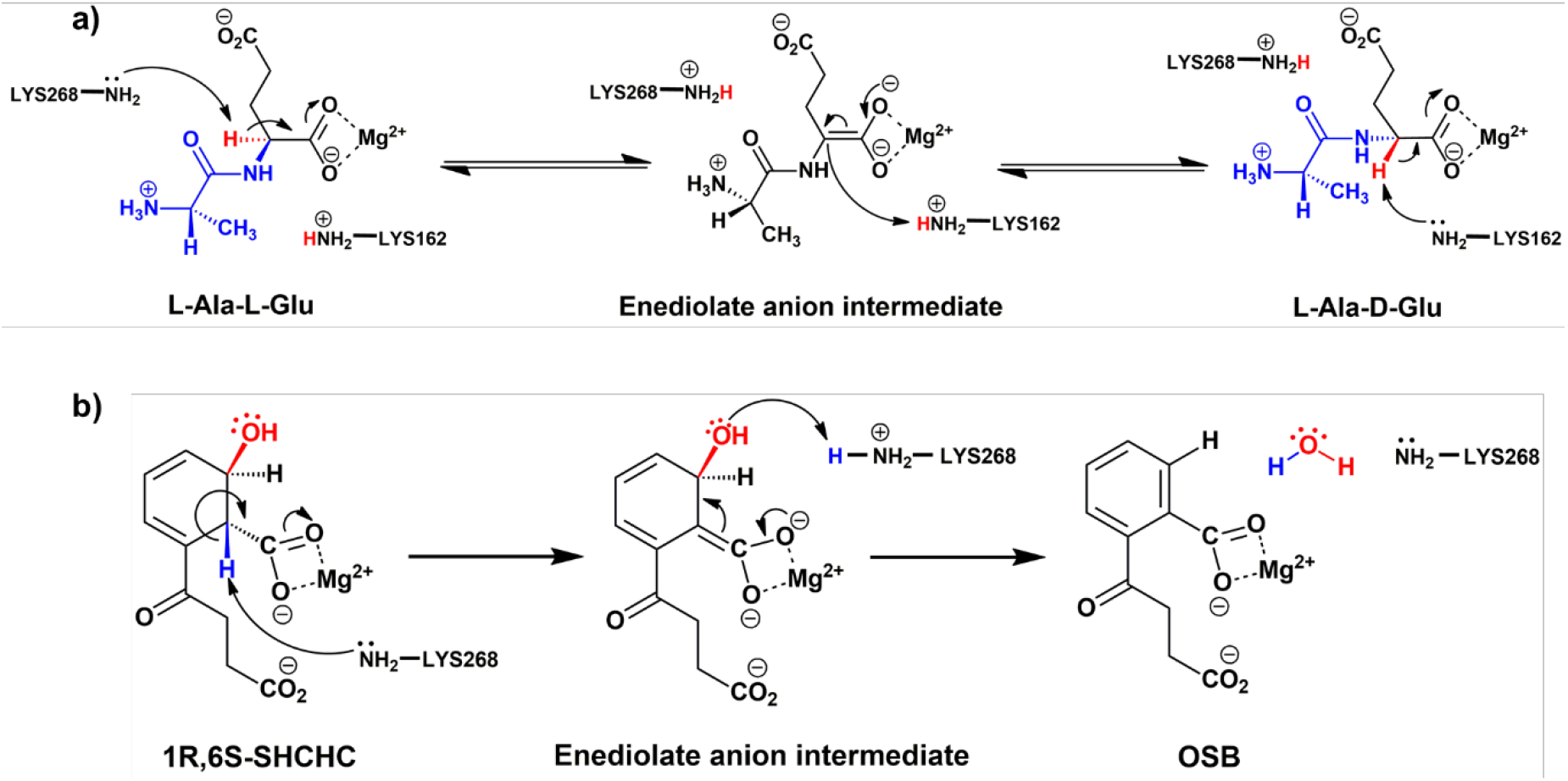
Proposed mechanism for a) AEE activity (1,1-proton transfer reaction) and b) OSBS activity (dehydration reaction)

In the present work, using comprehensive MD simulations supplemented by the hybrid QM/MM calculations, we have provided the mechanism of the spontaneous origin of promiscuity in the AEE enzyme, and we show how a mutation of Asp 321 to Glycine triggers an allosteric conformational change in the capping loop which benefits the substrate recognition for OSBS activity. In addition, our study provides a theoretical validation of the previously proposed reaction mechanism of AEE and OSBS activity for the first time.

## 2. Computational Method

In the present study, we have used Molecular Docking to get the initial structure of promiscuous substrate-bound complexes, MD simulations for the conformational sampling of the WT enzyme and its variant complexes, and hybrid QM/MM calculations for the potential energy surface (PES) scanning to get the reaction coordinate of the reactions. The details of each step are discussed below.

### 2.1. System setup

The initial coordinate for apo Ala-Glu epimerases and substrate-complexed Ala-Glu epimerases in *Bacillus subtilis* were taken from the protein data bank (PDB ID: 1JPM for Apo-and 1TKK for holo-enzyme).^25,26^ The coordinate of the non-native substrate, 2-succinyl-6-hydroxy-2,4-cyclohexadiene-1-carboxylate((1R,6S-SHCHC)) was taken from the already reported crystal structure of substrate-complexed OSBS (PDB ID: 1R6W)^27^. Molecular docking using Autodock Vina^28^ was used to get the Holoenzyme structure of Ala-Glu epimerase with the non-native substrate. The missing hydrogen and other heavy atoms were added using the LEAP module of the Amber20 package.^29^ The standard protonation state of the system was substantiated using the PropKa 3.0 program.^30^ The protein molecules and the native substrate L-Ala-L-Glu dipeptide were treated by Amber ff14SB forcefield. The parameters for non-standard substrate, 2-succinyl-6-hydroxy-2,4-cyclohexadiene-1-carboxylate((1R,6S-SHCHC)) were generated using the antechamber module of Amber20 for GAFF2 parameters. The partial atomic charges and missing parameters for both substrates were obtained from the RESP^31,32^ charge fitting method of a QM optimized geometry using the HF/6-31 g(d,p) level of theory. Since an Mg^2+^ metal ion was hexacoordinated with the enzyme as a cofactor, its parameters were prepared by the MCPB.py program, a python-based metal parameter builder used to generate a forcefield for the metal center of the protein, employing the bonded model approach.^33^ After proper parameterizations of the protein and substrate molecules, the systems were solvated in a truncated octahedral box with a TIP3P^34^ water model, extending up to 10 A° from the protein surface. The resulting charge of the prepared model was neutralized with counterions depending upon the total charge of the system.^35^

### 2.2. MD Simulation

All MD simulation was carried out using the Amber20 package in three different replicas starting from different initial velocities. The modeled systems were first minimized to remove the poor contacts and relax the system in two steps to get minimum energy conformation.^36^ In the first step, keeping the protein constrained by position restraint with a force constant of 500 Kcal/mol-Å^2^, and in the second step, it was a full minimization without any restraint. All minimization was done using 5000 steps of steepest descent^37^ and 5000 steps of conjugate gradient methods.^38^ Thereafter, systems were gently heated to the target temperature from 10 K to 300 K for a time period of 50 ps under constant volume periodic boundary conditions i.e. NVT ensemble. In the further step, the system is maintained at a constant temperature of 300 K and pressure of 1.0 atm in the NPT ensemble for 1 ns using the Langevin thermostat^39^ and Berendsen barostat^40^. Constant pressure was maintained with a relaxation time of 2 ps and temperature was controlled with a collision frequency of 1 ps^-1^. This was followed by equilibration for approximately 3 ns for each system which was further carried out to production simulation for 500 ns for each system. A cutoff of 10 A° was used with Particle Mesh Ewald (PME) method^41^ to treat long-range electrostatic and nonbonded interaction. During all MD simulations, all bonds involving hydrogen were constrained by the SHAKE^42^ algorithm.

All analyses of the trajectories were performed using the Cpptraj module, available in the Amber20 package. VMD^43^ and pyMol^44^ software is used for visualization purpose and rendering of the figure.

### 2.3. QM/MM Calculations

The reaction mechanism was studied for the representative frame of the most populated trajectories for each system by use of the QM/MM calculations. QM regions for WT AEE with native substrate included substrate (L-Ala-L-Glu), Mg^2+^ and ligated one water molecule and protein residues (D191, E219, D244), K162, K268, D321, and D323 while AEE with promiscuous substrate includes a substrate (1R,6S-SHCHC), Mg^2+^ and ligated one water molecule and protein residues (D191, E219, D244), K162, K268. The active region of QM/MM calculation includes all protein residues and water molecules within 8Å of the substrate. All QM/MM calculations were performed with ChemShell,^45,46^ by combining Turbomole^47^ for the QM part, and DL_POLY^48,49^ for the MM part. The MM region was described using the Amber ff14SB force field. The electronic embedding scheme was used to account for the polarizing effect of the enzyme environment on the QM region. The QM/MM boundary was treated using hydrogen link atoms with the charge-shift model.^50^ During QM/MM geometry optimizations, the QM region was treated using the hybrid UB3LYP^51^ functional with two basis sets, B1 and B2, where B1 stands for def2-SVP* and B2 is for def2-TZVP. For geometry optimization and frequency calculations, we used the B1 basis set. The energy was further corrected using B3LYP-D3^52^ functional. All of the QM/MM transition states (TSs) were located by relaxed potential energy surface (PES) scans followed by full TS optimizations using the dimer method. The energies were further corrected with the large all-electron def2-TZVP basis set. The zero-point energies (ZPEs) were calculated for all species, and the respective final energies are reported as UB3LYP/B2+ZPE data.

## 3. RESULTS AND DISCUSSION

### 3.1. MD Simulation of the substrate-free and substrate-bound dipeptide epimerase

The crystal structures of the substrate-free and substrate-bound (L-Ala-L-Glu dipeptide) enzyme show significant differences in the conformation of the capping loops in both forms; for instance, the capping loop is quite disordered in substrate-free form relative to the capping loop conformation of the substrate-bound enzyme. To study whether these conformational changes are substrate regulated, we, therefore, performed three separate MD simulations: a) of the substrate-free enzyme (Apo-form), b) substrate-docked in open cavity form of the enzyme and, c) substrate-bound co-crystal form of the enzymes.

The MD simulation of the substrate-free epimerase shows capping loop remains in the open conformation during the entire course of the simulations (see Figure 2a). We found an Arg 24 at the tip of the capping loop forming a salt bridge with Glu30 which is also part of the same loop. This salt bridge between Arg24-Glu30 blocks the movement of the capping loop towards the channel side and is the main driving force for the opening of the channel throughout the simulation. A distance plot between Arg24-Glu30 shows many fluctuations in the salt-bridge (Figure 2) and therefore, we believe a small perturbation in the channel side due to the entry of the substrate may disrupt the Arg24-Glu30 interactions.

**Figure 2.**
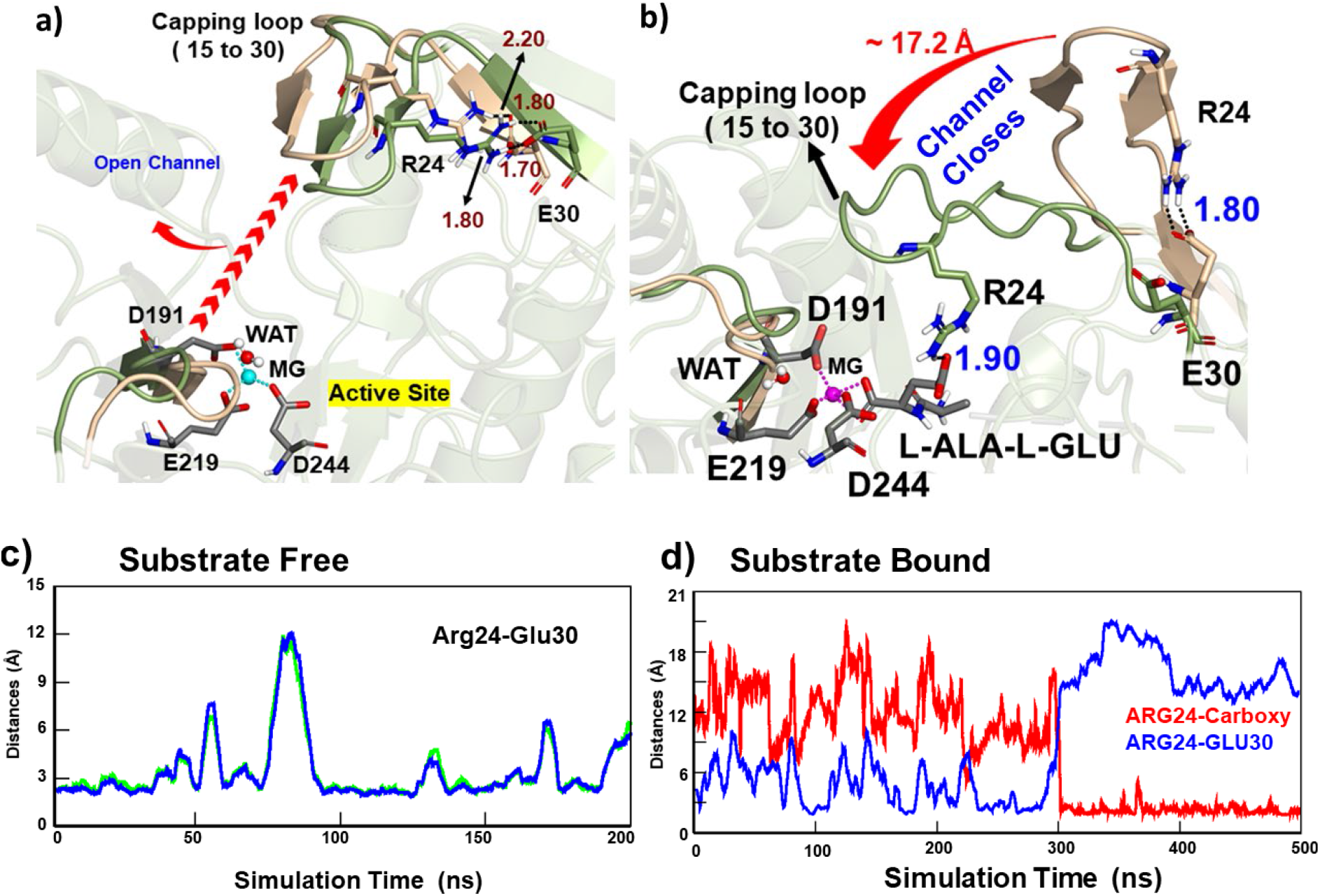
a) Superposition of initial and final snapshot during the simulation of the substrate-free enzyme (Apo form). The Colour of Arg24, Glu30, and the capping loop in the initial frame is shown in golden color while in the final frame they are in green color. Active site residues are in grey. b) Distance plot between Arg24 and Glu30 during the whole course of the simulation.

To further validate this conjecture, we docked the substrate in the open-channel state of the enzyme and performed MD simulations for 500 ns. As expected, during the simulations of the substrate-docked open-channel conformation, the Arg24-Glu30 salt bridge breaks and the capping loop moves towards the channel, which, consequently, closes the channel during the simulations (see Figures 2b and 2d). Interestingly, the Arg24 residue which initially formed a salt bridge with Glu30 changes its orientation towards the substrate after 300 ns of the simulations and now forms a strong hydrogen bond/salt-bridge from the carboxylate group of the L-Ala-L-Glu substrate. The interaction of the Arg24 with the substrate now tilts the loop towards the channel, and as a result, the channel closes. Here, we note that the experimental co-crystal of the enzyme with the same substrate (PDB id:1TKK) shows a closed-channel conformation where the capping loop forms a similar hydrogen bond with the substrate carboxylate group. In fact, the MD simulation of the co-crystallographic substrate-enzyme complex doesn’t show a change in the conformation of the capping loop (see SI for more details) and it remains closed for entire simulations (see figure S2). *Therefore, in nutshell, we conclude that the entry of the substrate transpires a huge conformational change in the enzyme and it acts as a signal to the enzyme for channel gating where the capping loop plays the role of gatekeeper*.

### 3.2. Mechanistic study of 1, 1-proton transfer reaction

The AEE enzyme is well known to catalyze the 1,1-proton transfer reaction to transform L-Ala-L-Glu dipeptide to L-Ala-D-Glu dipeptide as shown in Scheme 1a. To validate the mechanism, we performed the QM/MM calculations of a representative snapshot from the MD simulations of the enzyme-substrate complex. During the MD simulation, we observed that Lys162 and Lys268 are close to the reactive center of the substrate and can act as acid-base for the 1,1-proton transfer reaction (see Figure S3). To obtain the energy profile of the reaction, we performed the potential energy surface (PES) scanning of the QM/MM optimizes Michaelis Complex. As can be seen in the reaction profile, the first protonation starts as a proton transfer from Cα (see atom ‘a’ in Figure 3) carbon to the deprotonated Lys268 (K268) with a transition state barrier of 18.7 Kcal/mol and forms an enediolate anion intermediate IM. The intermediate IM is unstable and in the proceeding step it abstracts a proton from the other lysine, i.e., Lys162, in a very facile manner (TS barrier < 1 kcal/mol) to form a thermodynamically stable product (PC), L-Ala-D-Glu dipeptide, as shown in Figure 3. As can be seen, the first proton transfer from the Cα carbon of the substrate to K286 is the rate-determining step and the Mg^2+^ ion which is coordinated with the substrate provides stability to the transition state to lower the TS barrier of efficient reaction.

**Figure 3.**
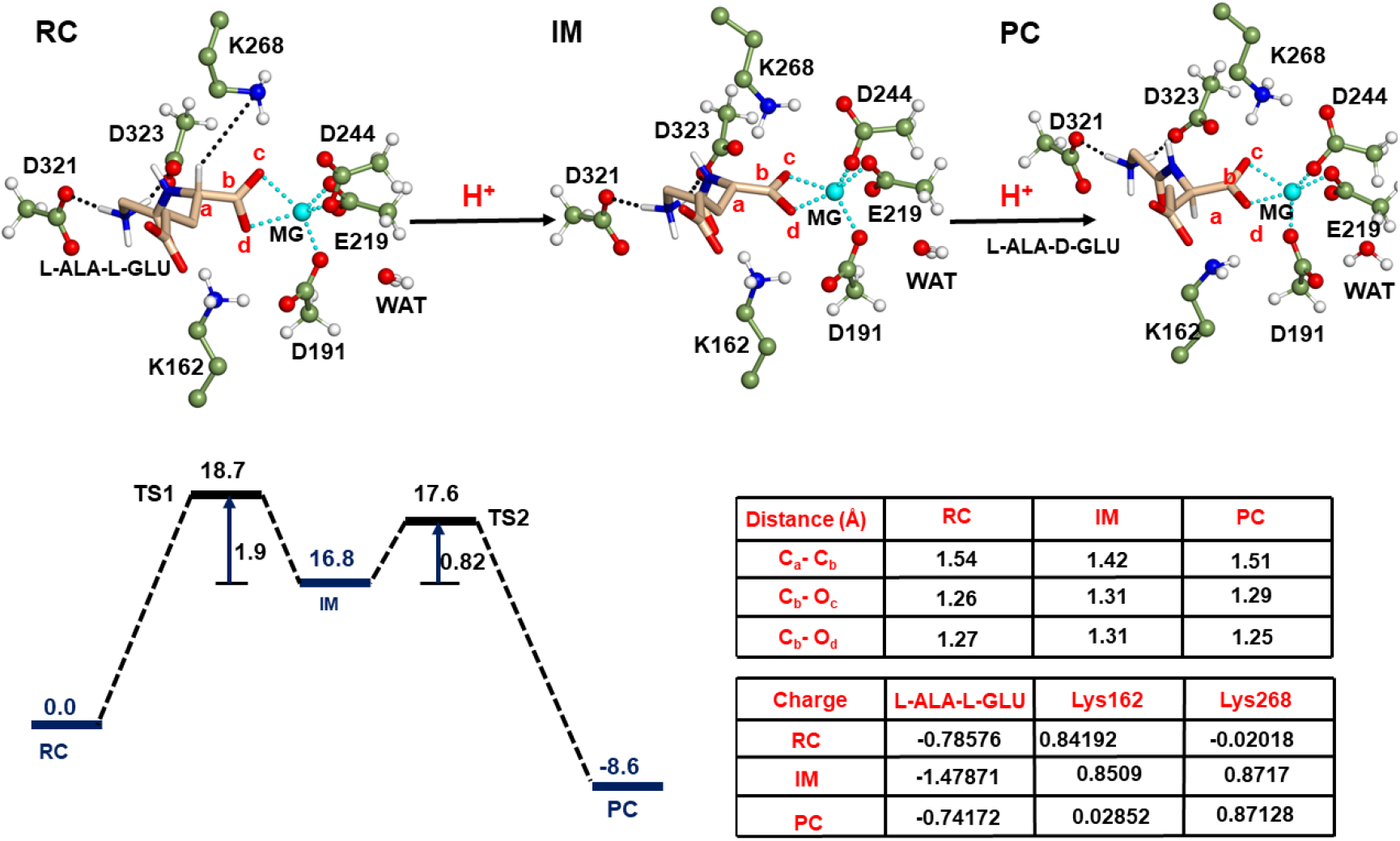
QM/MM/B3LYP-D3/def2-TZVP calculated mechanism for 1,1-Proton transfer reaction. Key residues of the QM region are shown by the ball and stick model. A corresponding energy profile and barriers for the process are shown in the schematic diagram at bottom of the figure. The reported energy values are in kcal/mol.

#### The allosteric effect on the Caping loop and its role in Catalysis

We have seen that the K162 and K268 residues act as acid-base pairs during the catalysis of the 1,1-proton transfer reactions. In addition, the entry of substrate instigates a large conformation change in the capping loop which doesn’t participate in the reaction itself. If so, then how does the capping loop come into play and why does Nature evolve it like a gatekeeper? To elaborate on the role of the capping loop we thoroughly studied the active site architecture of the AEE enzyme, and we found a well-organized active site engineering in the AEE enzyme. As can be seen from Figure 4, the α-ammonium group of the dipeptide substrate forms a hydrogen bond with Asp 321 and Asp 323 residues. In addition, the carboxylate group of the substrate forms a hydrogen bond with the guanidium group of the Arg24 which plays a major role in the opening and closing of the channel, and hence substrate can be a linker between the Asp 321 and Asp 323 residues. Therefore, *a small perturbation on one end of the active site (near D321-D333) can allosterically induce a large conformational change in the capping loop*.

**Figure 4.**
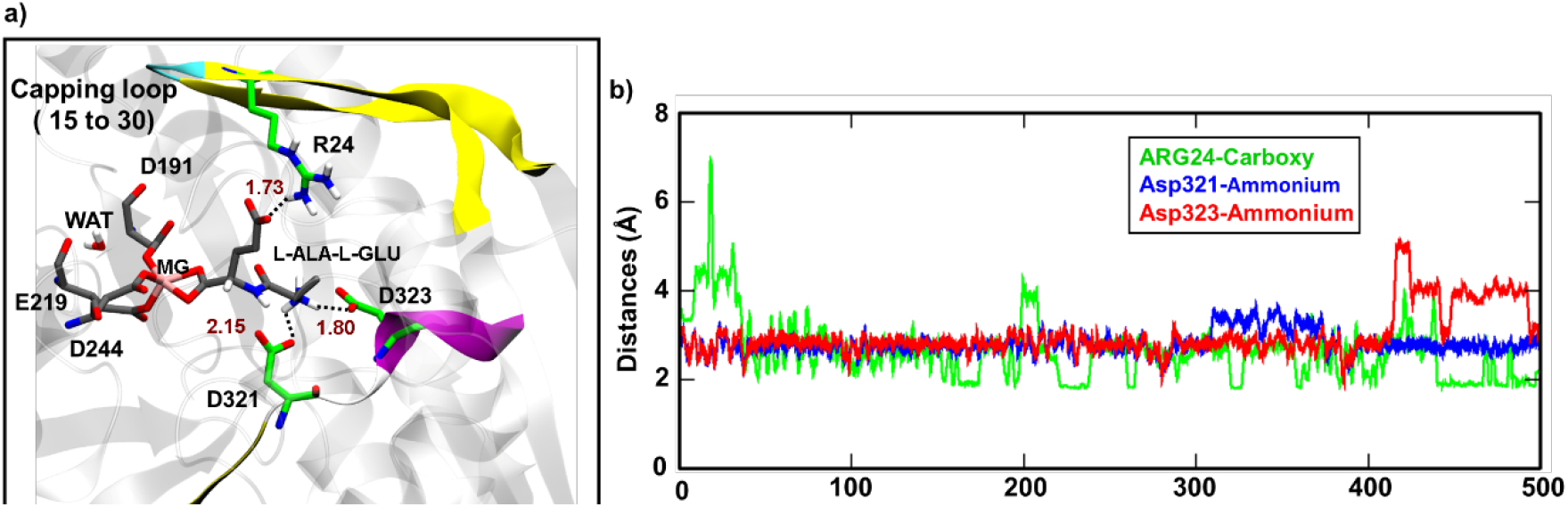
a) Snapshot showing the position of residue R24, D321, and D323; active site and substrate. b) Corresponding distance vs simulation time plot. Arg24 to carboxylate of the substrate (green), Asp321 to ammonium of substrate (blue), and Asp323 to ammonium of substrate (red).

To further substantiate the allostery effect, we mutated D321 which is far from the capping loop with Glycine and performed long time-scaled MD simulations for this mutant enzyme. To our surprise, we found that the capping loop which was initially interacting with the substrate through Arg24 and closing the channel of the enzyme breaks its interactions with the carboxylate (see Figure 5). Due to a loss of interaction with Arg24 and the carboxylate of the substrate, the capping loop restores its open-channel conformation in the D321G mutant. In this open-channel conformation of the D321G mutant, the substrate loses its interaction with Arg24 which weakens the binding of the native substrate and results in poor catalytic efficiency of the mutant towards the native substrate D-Ala-D-Glu. Our finding strongly correlated with the experimental finding by Schmidt et al where site-directed mutagenesis of D321G results in a complete loss of native activity of epimerase (cf. Figure 5c for comparative interaction energy of the substrate with the enzyme in WT and D321G mutant).

**Figure 5.**
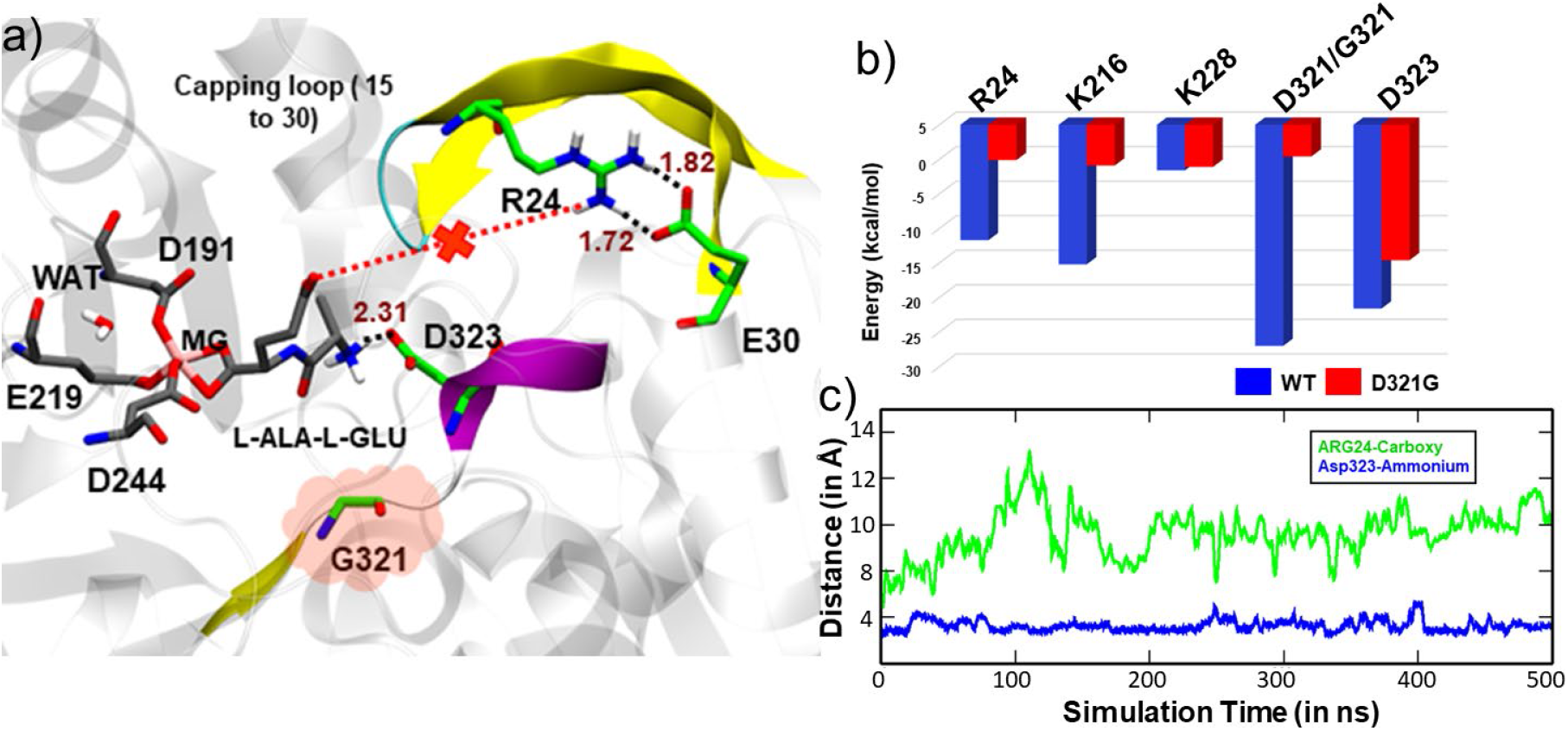
a) Snapshot showing the position of residue R24, E30, G321, and D323; active site and substrate. b) Corresponding distance vs simulation time plot. Arg24 to carboxylate of the substrate (green), Asp323 to ammonium of substrate (blue). c)Energy vs Residue plot of MMPBSA calculation. The blue bar is for WT AEE with native substrate and the red bar is for D321G AEE with the native substrate. The graph clearly shows how interaction energy gets reduced on mutation.

### 3.3 MD simulation of the D321G mutant with non-native substrate

We have seen that the D321G mutation weakens the interaction with the native substrate, opens the capping loop, and destroys the native activity. However, the study by Schmidt et al shows that the D321G mutation changes the AEE epimerase from a specialist to a promiscuous (i.e., 1,1-proton transfer of AEE to dehydration activity of OSBS) enzyme. To further elaborate on the spontaneous emergence of the promiscuity, we performed the MD simulations for the WT and D321G mutant with the non-native substrate, 1R,6S-SHCHC (RSH). In the WT simulation with substrate RSH, we found that D321 interacts with catalytic residue K162 and drags it away from the substrate which might be a reason for no activity for dehydration for RSH by the WT AEE enzyme (see figure 6). In contrast, in the MD simulation of the D321G mutant, due to the mutation of Asp to Gly, it loses the favorable interaction with Lys 162 and releases Lys 162 to interact with the carboxylate of the substrate which in turn increases the overall binding affinity of the substrate with the mutant AEE enzyme. Since in the mutant enzyme both catalytic Lysines (K162 and K268) are in close proximity to the substrate RSH, it may catalyze the dehydration activity of RSH and shows a promiscuous activity similar to OSBS enzyme (see figure S4).

**Figure 6.**
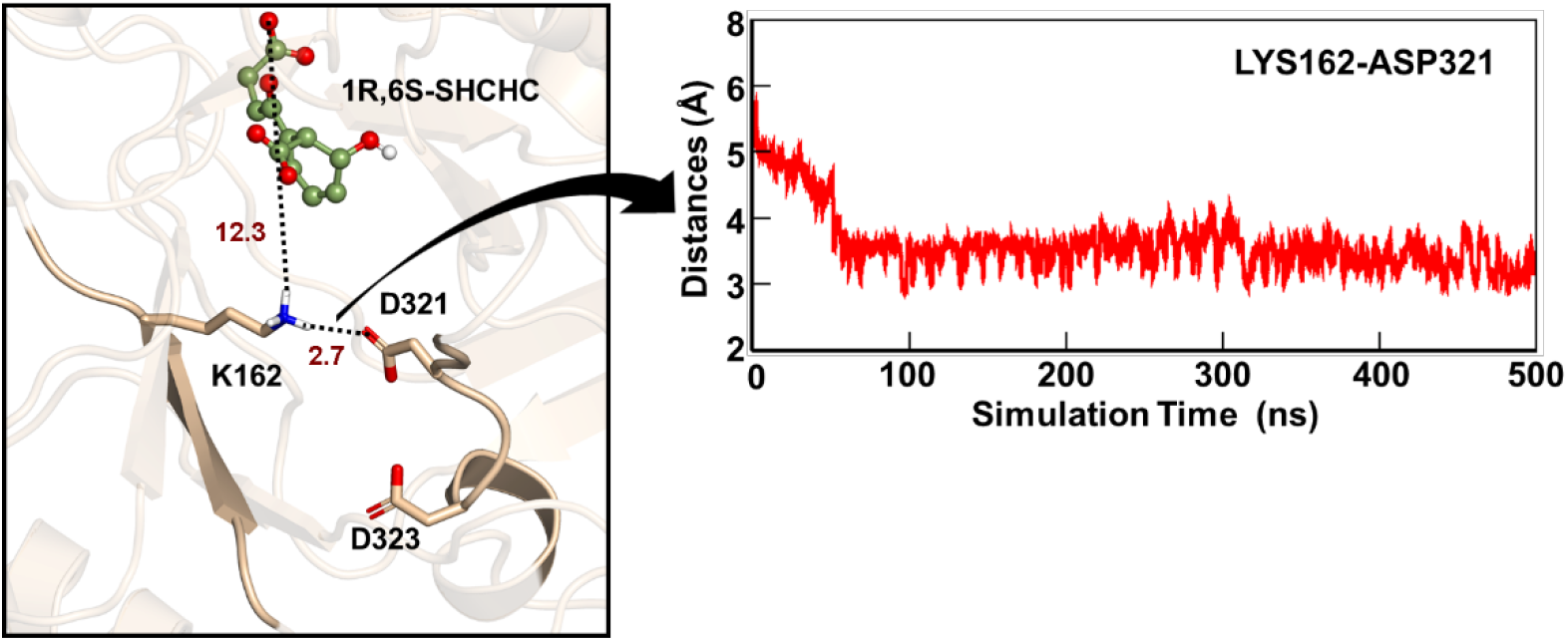
a) Most populated snapshot showing the position of residue K162, D321, D323, and substrate during the MD simulation of WT AEE with the non-native substrate (RSH). b) Corresponding distance vs simulation time plot of Asp321 and K162.

So far, we have seen that MD simulations of the D321G mutant with non-native substrate RSH provide a favorable condition for the secondary function for OSBS activity, however, it does not provide an insight into the mechanism of the reaction, we, therefore performed QM/MM calculations for a representative snapshot from the MD trajectory. The reaction coordinate for the proton abstraction from the Cα position of the substrate was obtained by potential energy surface (PES) scanning of a QM/MM optimized Michaelis Complex. As can be seen from the reaction profile in Figure 7, the reaction starts from the proton abstraction from the Cα (cf. atom **a** in Figure 7) carbon of the substrate to Lys268 with a small energy barrier of 14.0 kcal/mol and forms an enediolate intermediate IM. Here, other catalytic Lysine, i,e., Lys162 plays a crucial role in the stabilization of the transition state and lowers the transition state barrier. In the next step, the β-hydroxy end of endothermic IM easily (i.e., with a barrier of 6.8 kcal/mol) accepts one proton from the Lys162 and forms the product and a water molecule.

**Figure 7.**
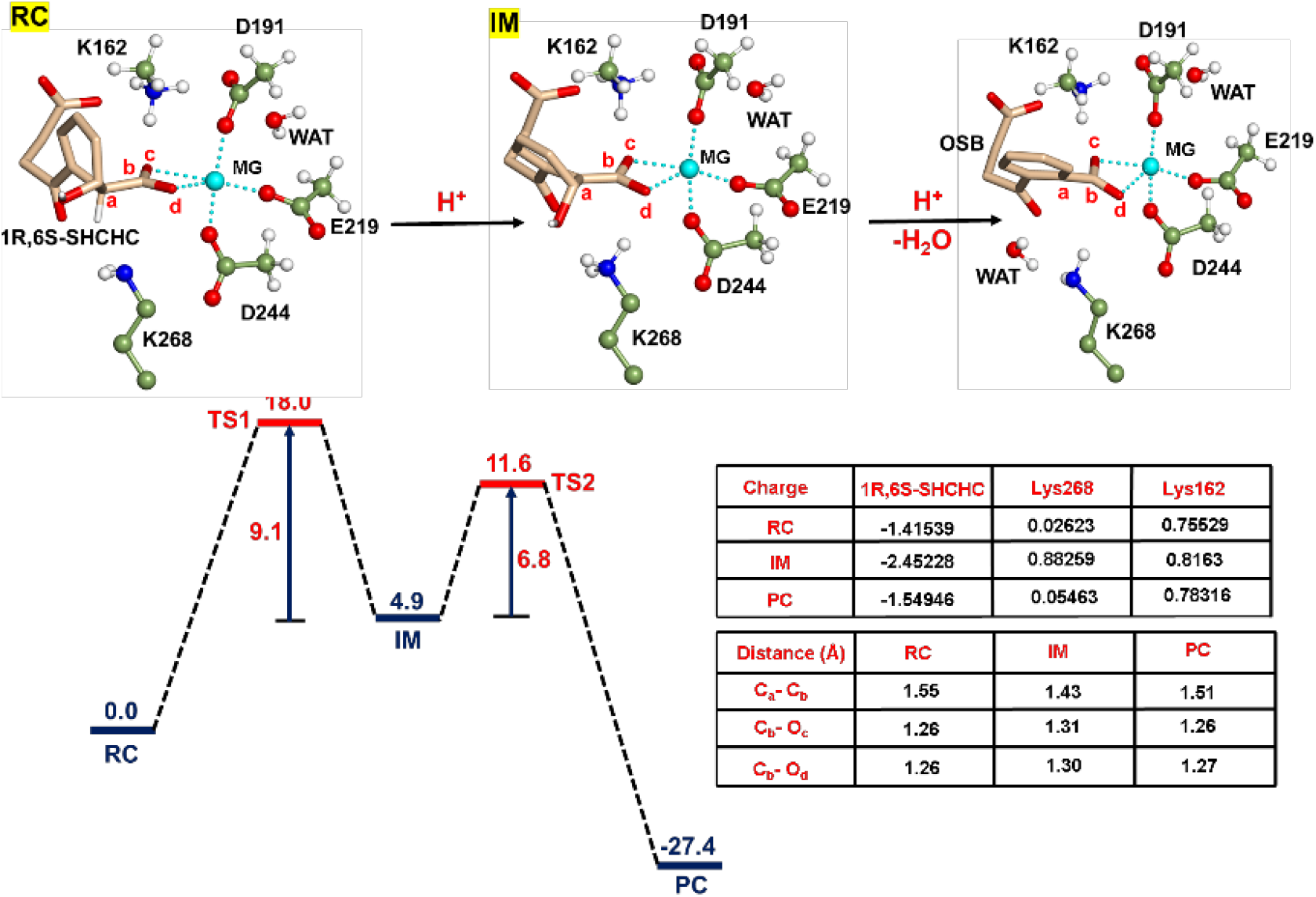
QM/MM/B3LYP-D3/def2-TZVP calculated mechanism for dehydration reaction. Key residues in the QM region are shown by the ball and stick model. A corresponding energy profile and barriers for the process are shown in the schematic diagram at bottom of the figure. The reported energy values are in kcal/mol.

In nutshell, the QM/MM calculations show that the conformation obtained from the MD simulations of the non-native substrate in the mutant enzyme can efficiently catalyze the secondary function and shows a significant degree of promiscuity.

## 4. Conclusion

Using extensive MD simulations and QM/MM calculations we have studied the mechanism of promiscuity due to a single site mutation in AEE epimerase. The role of the capping loop was well established in the substrate recognition but here, we have shown that the conformation of the capping loop is substrate driven and it is key to promiscuous functions. Using MD simulations of the WT and D321G mutant for both native and non-native substrates we have shown that the D321G mutation which is far from the capping loop causes an allosteric effect on the capping loop which further opens the channel for the non-native substrate. Our QM/MM calculations for the WT and mutant AEE epimerase provide a first-ever mechanistic validation of the 1,1-proton transfer and dehydration reaction by these enolase families. Since AEE epimerase possesses a highly conserved TIM-barrel fold, we believe that our study provides a crucial lead to understanding the mechanism of emergence of secondary function which can be useful to repurpose ancient enzymes for modern usage.

## 5. Acknowledgement

Authors acknowledge the Department of Biotechnology, Govt. of India for the Ramalingaswami Re-entry research grant (BT/RLS/Re-entry/2017/10) to support the resources needed for this project. The authors also acknowledge Prof. Sason Shaik, Department of Chemistry, The Hebrew University of Jerusalem Israel for partial assistance in computational work.

## Supporting Information

The supporting material contains, QM optimized geometries, different representative snapshots, etc.

