## Supporting Information for "Combined MD and QM/MM Calculations reveal Allostery Driven Promiscuity in Dipeptide Epimerases of Enolase Family"

**Table of Contents:**

**Figure S1**: a) Superposition of initial and final snapshot during the simulation of the substrate-complexed structure (PDB id. 1TKK). b) Distance plot between Arg24 and Glu30 during the whole course of simulation.

**Figure S2:** Representative snapshot from the most populated structure of the MD simulations showing the active site of WT AEE complexed with native substrate. b) Distance vs Simulation time plot between Lys162 and α-carbon of substrate, Lys268 and α-proton of substrate.

**Figure S3:** Representative snapshot from the most populated structure of the MD simulations showing the active site of D321G AEE complexed with non-native substrate. b) Distance vs Simulation time plot between Lys268 and α-proton of substrate, Lys268 and hydroxy group of substrate and Lys162 to carboxylate of substrate.

**Figure S4:** Representative snapshot from the most populated structure of the MD simulations showing the active site of D321GAEE complexed with the non-native substrate which clearly shows the close proximity of Lys 268 with Asp244 which supports the deprotonated state of Lys268. b) Distance vs Simulation time plot between Lys268 and Asp244.

**Figure S5:** Transition structure during PES for the 1,1-proton transfer in WT AEE complexed with native substrate.

**Figure S6:** Transition structure during PES for the dehydration in D321G AEE complexed with the non-native substrate.

- QM region coordinates of a QM/MM optimised geometry of WT AEE complexed with native substrate. (RC, TS1, IM, TS2 and PC)

- QM region of a QM/MM optimised geometry of D321G AEE complexed with the non-native substrate. (RC, TS1, IM, TS2 and PC)

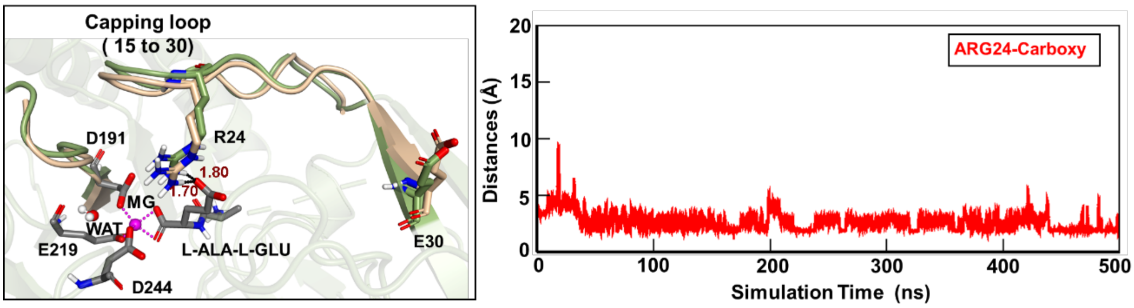


Figure S1. a) Superposition of initial and final snapshot during the simulation of the substrate-complexed structure (PDB id. 1TKK). The Colour of Arg24, Glu30 and the capping loop in the initial frame is shown in golden colour while in the final frame they are in green colour. Active site residues are in grey colour. b) Distance plot between Arg24 and Glu30 during the whole course of simulation.

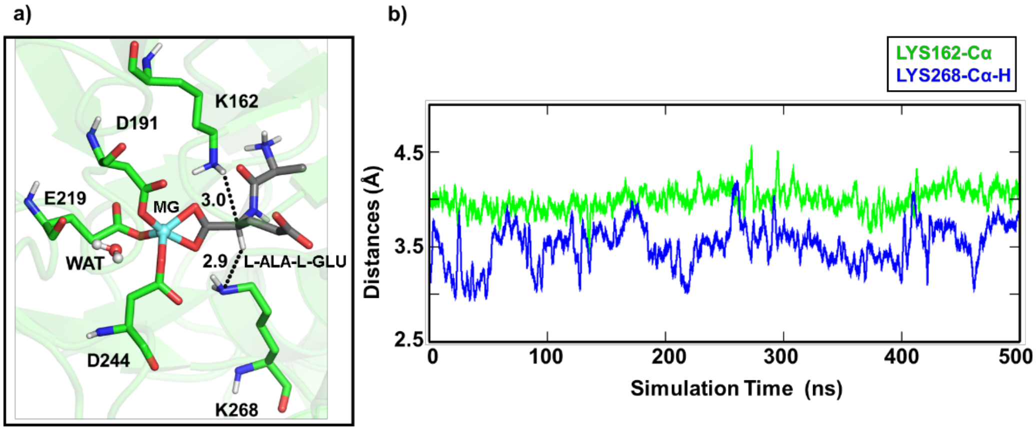


Figure S2. a) Representative snapshot from the most populated structure of the MD simulations showing the active site of WT AEE with native substrate b) Distance vs Simulation time plot between Lys162 and α-carbon (green), Lys268 and α-proton (blue). This snapshot clearly shows the close proximity of Lys pair with substrate.

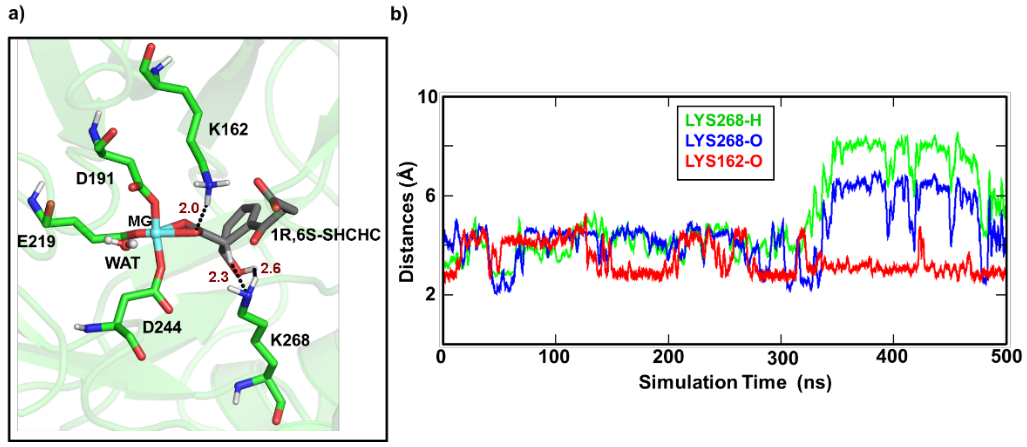


Figure S3. a) Representative snapshot from the most populated structure of the MD simulations showing the active site of D321G AEE with non-native substrate. b) Distance vs Simulation time plot between Lys268 and α-proton (green), Lys268 and hydroxy (blue) and Lys162 to carboxylate of substrate (red).

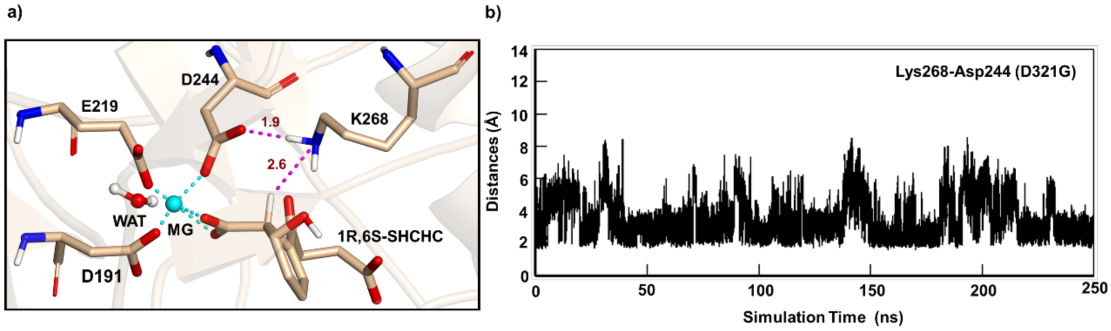


Figure S4. a) Representative snapshot from the most populated structure of the MD simulations showing the active site of D321GAEE with the non-native substrate which clearly shows the close proximity of Lys 268 with Asp244 which supports the deprotonated state of Lys268. b) Distance vs Simulation time plot between Lys268 and Asp244.

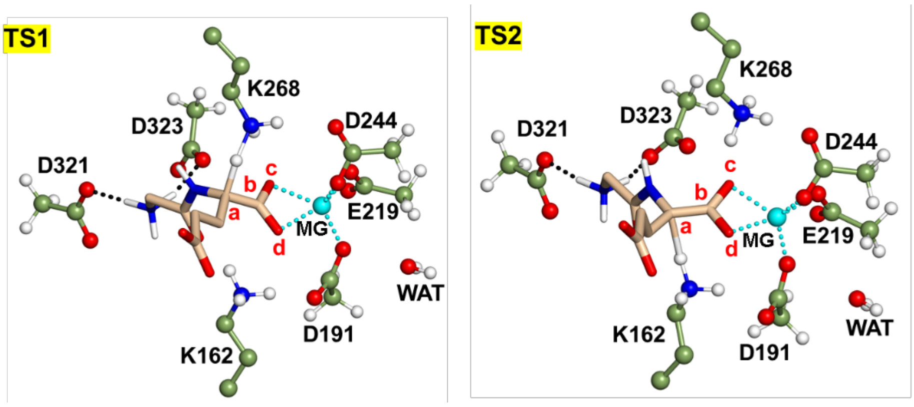


Figure S5. Transition structure during PES for the 1,1-proton transfer in WT AEE with native substrate.

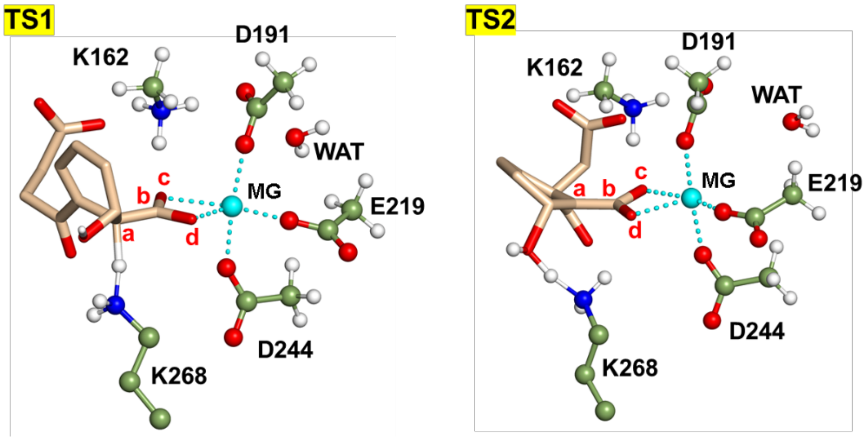


Figure S6. Transition structure during PES for the dehydration in D321G AEE with the non-native substrate.

QM region of a QM/MM optimised geometry of WT AEE with native substrate.

RC

| | | | |
|---|---|---|---|
| C | 46.1603109 | 29.6360920 | 38.3388999 |
| H | 46.7374204 | 30.5343737 | 38.0654182 |
| H | 45.8796023 | 29.1404682 | 37.3931735 |
| C | 44.8908490 | 30.0128402 | 39.1061394 |
| H | 44.2469228 | 29.1309909 | 39.2528771 |
| H | 45.0991409 | 30.3853929 | 40.1213364 |
| C | 44.0904048 | 31.0808957 | 38.3709691 |
| H | 44.6416344 | 32.0338547 | 38.3394078 |
| H | 43.9056055 | 30.7846239 | 37.3275575 |
| N | 42.7863672 | 31.3308522 | 39.0225196 |
| H | 42.1239217 | 30.5412642 | 38.8897652 |
| H | 42.8418761 | 31.3483115 | 40.0756591 |

| | | | |
|---|---|---|---|
| H | 42.3211580 | 32.1960534 | 38.6738958 |
| C | 44.5052544 | 32.3510298 | 42.4559410 |
| H | 44.6409961 | 33.4393942 | 42.5282641 |
| H | 45.0735857 | 31.9467622 | 41.6067257 |
| C | 43.0212770 | 32.0036250 | 42.3443602 |
| O | 42.6944684 | 30.9821785 | 41.6923294 |
| O | 42.2001776 | 32.7843846 | 42.9067018 |
| C | 41.6697062 | 34.7305203 | 45.3771501 |
| H | 41.1801681 | 33.7490856 | 45.3269068 |
| H | 41.0887647 | 35.3761556 | 46.0534668 |
| C | 41.6968957 | 35.3387591 | 43.9637469 |
| O | 40.6092978 | 35.1332116 | 43.3139875 |
| O | 42.6613145 | 35.9665667 | 43.5179776 |
| C | 38.0127540 | 33.5248410 | 45.1739505 |
| H | 38.6480016 | 34.4189253 | 45.0720004 |
| H | 38.5773274 | 32.8491361 | 45.8385961 |
| C | 37.9568037 | 32.8309623 | 43.8051034 |
| O | 36.8982978 | 32.5996518 | 43.2105219 |
| O | 39.1166534 | 32.4945091 | 43.3682897 |
| C | 34.2536521 | 37.2438538 | 39.3460442 |
| H | 34.7946899 | 38.0543689 | 38.8300209 |
| H | 34.4347324 | 37.3691840 | 40.4239089 |
| C | 34.7790525 | 35.8812321 | 38.9126184 |
| H | 34.2142764 | 35.0687169 | 39.4038130 |
| H | 34.6345098 | 35.7432797 | 37.8280689 |
| C | 36.2553711 | 35.6844575 | 39.2543697 |
| H | 36.5637733 | 34.6847237 | 38.9117038 |
| H | 36.8716945 | 36.4083188 | 38.6817038 |
| N | 36.5146777 | 35.7701447 | 40.6983937 |
| H | 37.4133134 | 35.3098309 | 40.8863815 |
| H | 36.6675665 | 36.7508145 | 40.9603281 |
| C | 41.7029837 | 39.6076927 | 37.8720800 |
| H | 42.1672213 | 39.9718662 | 38.8040669 |
| H | 40.6266497 | 39.4580944 | 38.0295855 |
| C | 42.4354573 | 38.3316367 | 37.4625049 |
| O | 43.6422224 | 38.4243853 | 37.1879763 |
| O | 41.7635987 | 37.2441219 | 37.4580951 |
| C | 44.5824219 | 38.0849651 | 31.9956711 |
| H | 45.2129842 | 37.4983248 | 31.3107907 |
| H | 45.2691376 | 38.7963064 | 32.4875436 |
| C | 44.1052516 | 37.1431549 | 33.1185351 |
| O | 42.8885616 | 37.0924857 | 33.4796052 |
| O | 45.0201068 | 36.4671299 | 33.6376498 |
| Mg | 40.4043384 | 33.4625985 | 42.2731945 |

| | | | |
|---|---|---|---|
| N | 42.5334384 | 35.5464876 | 35.6858410 |
| H | 42.4783487 | 36.2454968 | 36.5144102 |
| H | 42.7148905 | 36.1227169 | 34.7977635 |
| H | 43.3121950 | 34.8790862 | 35.8019797 |
| C | 41.2137358 | 34.8956644 | 35.6525169 |
| H | 40.4593340 | 35.6945051 | 35.6228752 |
| C | 41.0208446 | 34.0204493 | 36.9053165 |
| O | 41.9979994 | 33.4315521 | 37.3796006 |
| N | 39.7614231 | 33.9759654 | 37.3739582 |
| H | 39.0524903 | 34.4905663 | 36.8462998 |
| C | 39.1940370 | 33.2046930 | 38.4939796 |
| H | 38.1469125 | 33.5248419 | 38.5325710 |
| C | 39.2190431 | 31.6768348 | 38.3079364 |
| H | 40.2266289 | 31.3833181 | 37.9943357 |
| H | 39.0588964 | 31.2089987 | 39.2912833 |
| C | 38.1665152 | 31.1356130 | 37.3467613 |
| H | 37.1813056 | 31.5671597 | 37.5910697 |
| H | 38.3766845 | 31.4274770 | 36.3010386 |
| C | 37.9566060 | 29.5999967 | 37.3779134 |
| O | 36.9256977 | 29.1937926 | 36.7925513 |
| O | 38.7897300 | 28.8810444 | 37.9837334 |
| C | 39.7624560 | 33.5585474 | 39.8772899 |
| O | 38.9960406 | 34.0697691 | 40.7339935 |
| O | 40.9385573 | 33.2091752 | 40.2047544 |
| O | 40.9740854 | 30.6146987 | 45.3855359 |
| H | 41.7155111 | 31.1629070 | 45.7154494 |
| H | 40.7945693 | 30.0733039 | 46.1784106 |
| H | 46.8319282 | 28.9702240 | 38.8807644 |
| H | 44.9612347 | 31.9510726 | 43.3615808 |
| H | 42.6749838 | 34.6093835 | 45.7806471 |
| H | 37.0962645 | 33.8344836 | 45.6761811 |
| H | 33.1899653 | 37.3854714 | 39.1547802 |
| H | 41.8428889 | 40.4054092 | 37.1425993 |
| H | 43.8808168 | 38.7025626 | 31.4349653 |
| H | 41.1007383 | 34.3048719 | 34.7435268 |

TS1

| | | | |
|---|---|---|---|
| C | 46.0491467 | 29.6470988 | 38.3861729 |
| H | 46.6084296 | 30.5568752 | 38.1136259 |
| H | 45.7678529 | 29.1557970 | 37.4375769 |
| C | 44.7800584 | 30.0003806 | 39.1718361 |
| H | 44.1863466 | 29.0978790 | 39.3843053 |
| H | 45.0055539 | 30.4336610 | 40.1591555 |
| C | 43.9032720 | 30.9852222 | 38.4009163 |
| H | 44.4091817 | 31.9576955 | 38.2930886 |
| H | 43.7042502 | 30.6140068 | 37.3839103 |
| N | 42.5945712 | 31.2329311 | 39.0591425 |
| H | 41.9197643 | 30.4505175 | 38.9330804 |
| H | 42.6579388 | 31.2812146 | 40.1083657 |
| H | 42.1383273 | 32.0939428 | 38.6823868 |
| C | 44.4143500 | 32.3865804 | 42.5303008 |
| H | 44.5024067 | 33.4752147 | 42.6414075 |
| H | 44.9855810 | 32.0369919 | 41.6587055 |
| C | 42.9478625 | 31.9704680 | 42.4197251 |
| O | 42.6771337 | 30.9335997 | 41.7727319 |
| O | 42.0860627 | 32.7116272 | 42.9899772 |
| C | 41.6314577 | 34.7638295 | 45.4429664 |

| | | | |
|---|---|---|---|
| H | 41.1270110 | 33.7893911 | 45.4751186 |
| H | 41.0988607 | 35.4468914 | 46.1213924 |
| C | 41.5476410 | 35.2872405 | 43.9986723 |
| O | 40.3979445 | 35.0376039 | 43.4849009 |
| O | 42.4689741 | 35.8678681 | 43.4194035 |
| C | 37.8218658 | 33.8125415 | 44.9835779 |
| H | 38.4923087 | 34.6809824 | 45.0823405 |
| H | 38.3831865 | 32.9498331 | 45.3755624 |
| C | 37.5936254 | 33.6160579 | 43.4831460 |
| O | 36.6014125 | 34.1035270 | 42.9085146 |
| O | 38.4994823 | 32.9595363 | 42.8681862 |
| C | 34.1333409 | 36.8997086 | 38.5423554 |
| H | 34.1747314 | 36.4493135 | 37.5344866 |
| H | 34.6307414 | 37.8797718 | 38.4578166 |
| C | 34.9193358 | 36.0011317 | 39.5160386 |
| H | 34.8067333 | 36.3435544 | 40.5593608 |
| H | 34.5429050 | 34.9628259 | 39.4794026 |
| C | 36.4051929 | 36.0328024 | 39.1333149 |
| H | 36.4883719 | 35.8556306 | 38.0496311 |
| H | 36.7835940 | 37.0534847 | 39.3162356 |
| N | 37.3052279 | 35.0567767 | 39.7943236 |
| H | 38.2213846 | 35.5112905 | 39.9173613 |
| H | 36.9920844 | 34.8481008 | 40.7520769 |
| C | 41.6379563 | 39.6130624 | 37.9997186 |
| H | 42.0933877 | 40.0034519 | 38.9263652 |
| H | 40.5589409 | 39.5185739 | 38.1689276 |
| C | 42.3077550 | 38.2707542 | 37.7250213 |
| O | 43.4450766 | 38.2676252 | 37.2192934 |
| O | 41.6863885 | 37.2093292 | 38.0790676 |
| C | 44.5262578 | 37.9926590 | 32.0896814 |
| H | 45.1296129 | 37.3682075 | 31.4132939 |
| H | 45.2461593 | 38.6950969 | 32.5470547 |
| C | 44.0551779 | 37.1075059 | 33.2669368 |
| O | 42.8673244 | 37.1361771 | 33.7119146 |
| O | 44.9700089 | 36.4048744 | 33.7509238 |
| Mg | 40.3394951 | 33.4446852 | 42.2768581 |

| | | | |
|---|---|---|---|
| N | 42.4502645 | 35.6394607 | 36.0093028 |
| H | 42.4557431 | 36.2441675 | 36.8833938 |
| H | 42.6668014 | 36.2437513 | 35.1499066 |
| H | 43.1924991 | 34.9275983 | 36.0838529 |
| C | 41.1005947 | 35.0411300 | 35.9299198 |
| H | 40.3761607 | 35.8669293 | 35.9170069 |
| C | 40.8459172 | 34.1310956 | 37.1463648 |
| O | 41.8117695 | 33.5345609 | 37.6573272 |
| N | 39.5778030 | 34.0494191 | 37.5607272 |
| H | 38.8698572 | 34.5539741 | 37.0209021 |
| C | 39.0871621 | 33.2234023 | 38.6693167 |
| H | 37.9818114 | 33.9618826 | 39.1455763 |
| C | 38.9114530 | 31.7443236 | 38.3146280 |
| H | 39.8612654 | 31.3668114 | 37.9039750 |
| H | 38.7875871 | 31.2045567 | 39.2612006 |
| C | 37.7889186 | 31.3158338 | 37.3837970 |
| H | 36.8196706 | 31.7300482 | 37.7115723 |
| H | 37.9492949 | 31.6767847 | 36.3499083 |
| C | 37.5918222 | 29.7787028 | 37.3076673 |
| O | 36.5078046 | 29.3975757 | 36.8031502 |
| O | 38.4961568 | 29.0229045 | 37.7489729 |
| C | 39.9392563 | 33.3740813 | 39.8962839 |
| O | 40.1746765 | 34.5218386 | 40.4067968 |
| O | 40.2917270 | 32.3371736 | 40.5459675 |
| O | 40.9350613 | 30.6858875 | 45.2959943 |
| H | 41.6867370 | 31.2248099 | 45.6173622 |
| H | 40.7427264 | 30.1423320 | 46.0817559 |
| H | 46.7433249 | 28.9854768 | 38.9042849 |
| H | 44.9055790 | 31.9754901 | 43.4122113 |
| H | 42.6551806 | 34.6404138 | 45.7962751 |
| H | 36.9495368 | 34.0068106 | 45.6075721 |
| H | 33.0775722 | 37.0851355 | 38.7399323 |
| H | 41.8146048 | 40.3727320 | 37.2382959 |
| H | 43.8450663 | 38.6260342 | 31.5214550 |
| H | 40.9904350 | 34.4912744 | 34.9952590 |

IM

| | | | |
|---|---|---|---|
| C | 46.0084984 | 29.6974068 | 38.4133187 |
| H | 46.5792687 | 30.6069969 | 38.1622636 |
| H | 45.7346718 | 29.2295121 | 37.4504824 |
| C | 44.7279436 | 30.0532466 | 39.1848405 |
| H | 44.0841150 | 29.1679681 | 39.3042348 |
| H | 44.9313682 | 30.3985492 | 40.2102890 |
| C | 43.9342717 | 31.1353008 | 38.4502535 |
| H | 44.4766776 | 32.0937581 | 38.4639266 |
| H | 43.8058509 | 30.8613737 | 37.3925614 |
| N | 42.5821180 | 31.3803364 | 39.0175138 |
| H | 41.9324602 | 30.5853954 | 38.8492953 |
| H | 42.5788283 | 31.4058950 | 40.0673845 |
| H | 42.1550719 | 32.2450122 | 38.5998857 |
| C | 44.3608381 | 32.3812228 | 42.5165660 |
| H | 44.4254119 | 33.4716973 | 42.6311835 |
| H | 44.9414206 | 32.0494313 | 41.6438313 |
| C | 42.8968565 | 31.9447380 | 42.3859691 |
| O | 42.6428240 | 30.9101918 | 41.7299344 |
| O | 42.0201935 | 32.6814311 | 42.9369531 |
| C | 41.6194407 | 34.6962344 | 45.4231516 |
| H | 41.1444893 | 33.7065446 | 45.4509195 |
| H | 41.0590631 | 35.3661254 | 46.0929867 |
| C | 41.5319386 | 35.2119820 | 43.9742269 |
| O | 40.3825679 | 34.9619875 | 43.4697146 |
| O | 42.4523743 | 35.7973048 | 43.3944601 |
| C | 37.7911898 | 33.9057612 | 45.0647374 |
| H | 38.4305404 | 34.7939566 | 45.1885631 |
| H | 38.3624315 | 33.0577095 | 45.4711637 |
| C | 37.6334416 | 33.7069109 | 43.5579741 |
| O | 36.7421582 | 34.3213621 | 42.9092155 |

| | | | |
|---|---|---|---|
| O | 38.4758192 | 32.9432666 | 43.0172822 |
| C | 34.0422832 | 36.7614058 | 38.5551832 |
| H | 33.9602340 | 35.9040544 | 37.8646111 |
| H | 34.4583473 | 37.5843179 | 37.9545952 |
| C | 35.0046752 | 36.3864872 | 39.7109719 |
| H | 35.5858658 | 37.2692832 | 40.0272064 |
| H | 34.4671260 | 36.0373506 | 40.6071241 |
| C | 35.9496177 | 35.2737002 | 39.2227943 |
| H | 35.4175147 | 34.3116148 | 39.1655694 |
| H | 36.3076045 | 35.5101437 | 38.2132572 |
| N | 37.1858395 | 35.0306011 | 40.0200933 |
| H | 37.8234284 | 35.8463777 | 39.9700477 |
| H | 37.0197786 | 34.7885601 | 41.0240508 |
| C | 41.5587328 | 39.6336401 | 38.0547390 |
| H | 42.1497913 | 39.9425161 | 38.9354925 |
| H | 40.5038563 | 39.7328624 | 38.3403736 |
| C | 41.8931291 | 38.1679986 | 37.7992597 |
| O | 42.8882127 | 37.9064382 | 37.0543859 |
| O | 41.2030683 | 37.3017444 | 38.3863596 |
| C | 44.5962663 | 38.0540166 | 31.9846882 |
| H | 45.2013765 | 37.4552381 | 31.2862823 |
| H | 45.3061710 | 38.7506303 | 32.4623048 |
| C | 44.1132243 | 37.1244909 | 33.1154510 |
| O | 42.8825099 | 36.9990872 | 33.3927702 |
| O | 45.0355737 | 36.5305795 | 33.7199300 |
| Mg | 40.2564917 | 33.3903349 | 42.1930932 |
| N | 42.5345931 | 35.6691178 | 35.7533053 |
| H | 42.7267806 | 36.4812550 | 36.4470298 |
| H | 42.7213006 | 36.1023276 | 34.7975373 |
| H | 43.2152639 | 34.9119838 | 35.9172837 |
| C | 41.1465368 | 35.1864979 | 35.8586208 |
| H | 40.4820239 | 36.0580693 | 35.9359712 |
| C | 40.9718849 | 34.2692121 | 37.0766338 |
| O | 41.9691263 | 33.6536223 | 37.5002466 |
| N | 39.7257614 | 34.1444579 | 37.5440656 |
| H | 38.9884905 | 34.6504322 | 37.0469443 |
| C | 39.3455861 | 33.1198264 | 38.4918463 |
| H | 37.7461328 | 34.2438306 | 39.5786866 |
| C | 39.2604994 | 31.7163409 | 37.9188403 |
| H | 40.0464362 | 31.6000241 | 37.1457335 |
| H | 39.5170060 | 31.0410826 | 38.7387919 |
| C | 37.9515690 | 31.1745769 | 37.3419455 |
| H | 37.0901434 | 31.5787063 | 37.9048208 |
| H | 37.7954488 | 31.4731936 | 36.2900057 |
| C | 37.7909912 | 29.6329061 | 37.3903254 |
| O | 36.7426934 | 29.1862816 | 36.8573895 |
| O | 38.6601401 | 28.9298568 | 37.9675504 |
| C | 39.8454596 | 33.2539872 | 39.8090094 |
| O | 40.1423661 | 34.3944663 | 40.3804841 |
| O | 39.9674808 | 32.1990835 | 40.5753382 |
| O | 40.9685499 | 30.6791996 | 45.3521225 |
| H | 41.7253883 | 31.2149189 | 45.6648932 |
| H | 40.7884917 | 30.1299908 | 46.1379093 |
| H | 46.6948479 | 29.0196515 | 38.9209005 |
| H | 44.8655150 | 31.9784240 | 43.3947013 |
| H | 42.6419826 | 34.6082129 | 45.7902041 |
| H | 36.8990883 | 34.0717499 | 45.6686163 |
| H | 33.0075458 | 37.0300619 | 38.7678087 |
| H | 41.7819423 | 40.3617030 | 37.2748869 |
| H | 43.8909026 | 38.6828601 | 31.4414709 |
| H | 40.8831930 | 34.6462036 | 34.9493358 |

TS2

| | | | |
|---|---|---|---|
| C | 45.8717826 | 29.9389073 | 38.6035548 |
| H | 46.4123731 | 30.8955878 | 38.5105914 |
| H | 45.6391438 | 29.6219538 | 37.5703920 |
| C | 44.5519771 | 30.1275392 | 39.3779529 |
| H | 43.9178352 | 29.2350777 | 39.2557797 |
| H | 44.7045555 | 30.2163817 | 40.4648164 |
| C | 43.7486753 | 31.3451884 | 38.8881609 |
| H | 44.1925886 | 32.2842039 | 39.2536817 |
| H | 43.7621115 | 31.4298481 | 37.7944590 |
| N | 42.3284762 | 31.3210871 | 39.3143731 |
| H | 41.8801763 | 30.4035503 | 39.1362378 |
| H | 42.2159377 | 31.3582160 | 40.3407652 |
| H | 41.6330817 | 32.0819374 | 38.8522922 |
| C | 44.3106521 | 32.3217292 | 42.5619928 |
| H | 44.3782375 | 33.4150512 | 42.6289406 |
| H | 44.8613605 | 31.9634052 | 41.6794900 |
| C | 42.8404171 | 31.8783297 | 42.4706863 |
| O | 42.5786263 | 30.6919076 | 42.1944269 |
| O | 41.9700949 | 32.7950392 | 42.6275897 |
| C | 41.5817485 | 34.5946613 | 45.3450502 |
| H | 41.1551641 | 33.5819140 | 45.3593528 |
| H | 40.9776850 | 35.2276636 | 46.0141793 |
| C | 41.4800689 | 35.1351421 | 43.8979745 |
| O | 40.3344454 | 34.8849519 | 43.3769587 |
| O | 42.3889176 | 35.7615540 | 43.3458570 |
| C | 37.8020493 | 33.9109671 | 45.1417720 |
| H | 38.4294572 | 34.8075325 | 45.2632809 |
| H | 38.3736257 | 33.0645436 | 45.5494547 |
| C | 37.6184829 | 33.7157136 | 43.6415682 |
| O | 36.7804495 | 34.4148478 | 43.0071935 |
| O | 38.3742374 | 32.8722108 | 43.0898762 |
| C | 34.1057842 | 36.7806324 | 38.6035505 |
| H | 34.0638127 | 35.8755516 | 37.9724191 |
| H | 34.5199067 | 37.5745239 | 37.9633115 |
| C | 35.0349541 | 36.5101230 | 39.8064202 |
| H | 35.4315382 | 37.4604368 | 40.2036019 |
| H | 34.4934613 | 36.0277284 | 40.6370463 |
| C | 36.2066829 | 35.5994646 | 39.3945446 |
| H | 35.8627987 | 34.5698088 | 39.2138790 |
| H | 36.6763169 | 35.9618006 | 38.4690478 |
| N | 37.2943257 | 35.5238365 | 40.4084331 |
| H | 37.6923431 | 36.4634589 | 40.5644393 |
| H | 36.9967522 | 35.1353740 | 41.3330934 |
| C | 41.6346923 | 39.6325026 | 38.0160437 |
| H | 42.2212469 | 39.9573487 | 38.8942028 |
| H | 40.5792294 | 39.6922265 | 38.3098713 |

| | | | |
|---|---|---|---|
| C | 42.0295966 | 38.1829970 | 37.7455154 |
| O | 43.0095650 | 37.9731226 | 36.9723817 |
| O | 41.3951076 | 37.2852380 | 38.3518519 |
| C | 44.6368063 | 38.1542996 | 31.9130254 |
| H | 45.2501727 | 37.5724207 | 31.2089081 |
| H | 45.3302805 | 38.8573477 | 32.4045667 |
| C | 44.1569980 | 37.1999998 | 33.0195009 |
| O | 42.9209556 | 37.0691462 | 33.2774265 |
| O | 45.0722275 | 36.5941112 | 33.6212664 |
| Mg | 40.0782415 | 33.2857707 | 42.1111827 |
| N | 42.6353586 | 35.7498051 | 35.5940275 |
| H | 42.8334083 | 36.5293488 | 36.3033919 |
| H | 42.7764780 | 36.2066042 | 34.6402085 |
| H | 43.3421739 | 35.0040653 | 35.7248486 |
| C | 41.2815557 | 35.1986319 | 35.7500838 |
| H | 40.5811159 | 36.0138837 | 35.9784935 |
| C | 41.2982121 | 34.1618264 | 36.8768895 |
| O | 42.3717532 | 33.6234955 | 37.1690039 |
| N | 40.1165020 | 33.8573600 | 37.4319177 |
| H | 39.2807567 | 34.3561076 | 37.1125211 |
| C | 40.0143287 | 32.7377170 | 38.3247083 |
| H | 38.1524770 | 34.9439554 | 40.1292501 |
| C | 39.7067889 | 31.3984396 | 37.6867464 |
| H | 40.2753846 | 31.3199773 | 36.7428422 |
| H | 40.0694839 | 30.6090473 | 38.3407428 |
| C | 38.2499085 | 30.9886096 | 37.4314022 |
| H | 37.5875207 | 31.4595240 | 38.1812695 |
| H | 37.8798624 | 31.3133547 | 36.4457314 |
| C | 37.9742907 | 29.4663760 | 37.5436646 |
| O | 36.8994625 | 29.0743261 | 37.0218948 |
| O | 38.7850235 | 28.7351355 | 38.1705216 |
| C | 39.7930602 | 32.9815449 | 39.6870534 |
| O | 39.7023037 | 34.1846316 | 40.2344931 |
| O | 39.8071360 | 31.9942883 | 40.5393085 |
| O | 40.9368281 | 30.6696835 | 45.4252974 |
| H | 41.7259208 | 31.1589540 | 45.7373737 |
| H | 40.7591884 | 30.0978758 | 46.1964890 |
| H | 46.5691170 | 29.2088251 | 39.0143733 |
| H | 44.8518681 | 31.9376206 | 43.4266390 |
| H | 42.5976776 | 34.5566462 | 45.7381280 |
| H | 36.9021900 | 34.0750545 | 45.7345584 |
| H | 33.0633080 | 37.0432678 | 38.7833648 |
| H | 41.8216334 | 40.3748731 | 37.2401531 |
| H | 43.9148070 | 38.7708567 | 31.3776362 |
| H | 40.9668909 | 34.7281634 | 34.8185729 |

PC

| | | | |
|---|---|---|---|
| C | 46.0357834 | 29.5857684 | 38.4127980 |
| H | 46.6100399 | 30.3974084 | 37.9354570 |
| H | 45.5872827 | 28.9981945 | 37.5903693 |
| C | 44.9365336 | 30.1589348 | 39.3164932 |
| H | 44.4789721 | 29.3719258 | 39.9407334 |
| H | 45.3684428 | 30.8905531 | 40.0230721 |
| C | 43.8056898 | 30.8489592 | 38.5445617 |
| H | 44.2172532 | 31.6826170 | 37.9449214 |
| H | 43.4050208 | 30.1240930 | 37.8121633 |
| N | 42.6948196 | 31.3445791 | 39.3306196 |
| H | 42.1361789 | 30.6039844 | 39.7733928 |
| H | 42.9686240 | 32.0066612 | 40.0520472 |
| H | 41.2057757 | 32.5290246 | 38.3931153 |
| C | 44.1900354 | 32.2277633 | 42.6404936 |
| H | 44.3255732 | 33.3166490 | 42.5910020 |
| H | 44.5883257 | 31.7847977 | 41.7151503 |
| C | 42.6871634 | 31.9259122 | 42.7329634 |
| O | 42.2583219 | 30.9319432 | 43.3616055 |
| O | 41.9402392 | 32.7530024 | 42.1155370 |
| C | 41.5954067 | 34.5283335 | 45.3326337 |
| H | 41.1670029 | 33.5140530 | 45.3647680 |
| H | 40.9889910 | 35.1737065 | 45.9877702 |
| C | 41.5118217 | 35.0514694 | 43.8831533 |
| O | 40.3610439 | 34.8171974 | 43.3534969 |
| O | 42.4314761 | 35.6415139 | 43.3115401 |
| C | 37.8174806 | 33.9008581 | 45.0609051 |
| H | 38.4890236 | 34.7489115 | 45.2661092 |
| H | 38.3607524 | 32.9882870 | 45.3448045 |
| C | 37.5732985 | 33.9236322 | 43.5568484 |
| O | 36.8153069 | 34.7883511 | 43.0574067 |
| O | 38.2130654 | 33.0805682 | 42.8503421 |
| C | 34.0386699 | 36.7789841 | 38.6017594 |
| H | 33.9888926 | 35.8449166 | 38.0167183 |
| H | 34.4465394 | 37.5418735 | 37.9209932 |
| C | 34.9677433 | 36.5720341 | 39.8125308 |
| H | 35.3902607 | 37.5368720 | 40.1412613 |
| H | 34.4265045 | 36.1606366 | 40.6807045 |
| C | 36.1103443 | 35.6152307 | 39.4493734 |
| H | 35.7508788 | 34.5792859 | 39.3526271 |
| H | 36.5646700 | 35.8997128 | 38.4887027 |
| N | 37.2105243 | 35.6032582 | 40.4462067 |
| H | 37.6202725 | 36.5482249 | 40.5345808 |
| H | 36.9456622 | 35.2605646 | 41.4143412 |
| C | 41.6349439 | 39.6244657 | 38.0086998 |
| H | 42.2285562 | 39.9376794 | 38.8862501 |
| H | 40.5814851 | 39.6865167 | 38.3103356 |
| C | 42.0189469 | 38.1750343 | 37.7193531 |
| O | 42.9686382 | 37.9702240 | 36.9079542 |
| O | 41.4037100 | 37.2742262 | 38.3414286 |
| C | 44.6514108 | 38.1967733 | 31.8801299 |
| H | 45.2644667 | 37.6203362 | 31.1717240 |
| H | 45.3413900 | 38.8958015 | 32.3814032 |
| C | 44.1613008 | 37.2294755 | 32.9677640 |
| O | 42.9164487 | 37.0838594 | 33.1830714 |
| O | 45.0623689 | 36.6272523 | 33.5917703 |
| Mg | 40.0216227 | 33.3377399 | 42.0320284 |
| N | 42.5738013 | 35.8057836 | 35.4674701 |
| H | 42.7830281 | 36.5771491 | 36.1850142 |
| H | 42.7316322 | 36.2560179 | 34.5101616 |
| H | 43.2636257 | 35.0473763 | 35.6149722 |
| C | 41.2120727 | 35.2792483 | 35.6289573 |
| H | 40.5215828 | 36.1005962 | 35.8668037 |

| | | | |
|---|---|---|---|
| C | 41.2572214 | 34.2404117 | 36.7493955 |
| O | 42.3476596 | 33.7575077 | 37.0419515 |
| N | 40.0900845 | 33.8553610 | 37.3091732 |
| H | 39.2074987 | 34.2895673 | 37.0324548 |
| C | 40.1231313 | 32.7235442 | 38.2133999 |
| H | 38.0254047 | 35.0102868 | 40.1497493 |
| C | 39.6489150 | 31.3898421 | 37.6286620 |
| H | 40.1097761 | 31.3063047 | 36.6310480 |
| H | 40.1042624 | 30.6240279 | 38.2605438 |
| C | 38.1663382 | 31.0399284 | 37.5373232 |
| H | 37.5823429 | 31.5781081 | 38.3060907 |
| H | 37.7274374 | 31.3022639 | 36.5636642 |
| C | 37.8839141 | 29.5359563 | 37.7773380 |
| O | 36.9059397 | 29.0450354 | 37.1675548 |
| O | 38.6186221 | 28.9167583 | 38.5994690 |
| C | 39.7017806 | 33.0084566 | 39.6346753 |
| O | 39.6429291 | 34.2104710 | 40.0934754 |
| O | 39.6027689 | 32.0379820 | 40.4228034 |
| O | 40.4004691 | 31.2968543 | 45.2470593 |
| H | 41.1487414 | 31.2137600 | 44.6071521 |
| H | 40.7781167 | 30.9515426 | 46.0708743 |
| H | 46.7469777 | 28.9378281 | 38.9250901 |
| H | 44.8025460 | 31.8725857 | 43.4691955 |
| H | 42.6099281 | 34.4927899 | 45.7295594 |
| H | 36.9393866 | 34.0557383 | 45.6878263 |
| H | 32.9996196 | 37.0509088 | 38.7874973 |
| H | 41.8201481 | 40.3721769 | 37.2375355 |
| H | 43.9246970 | 38.8122557 | 31.3499077 |
| H | 40.8819196 | 34.8078842 | 34.7032780 |

QM region of a QM/MM optimised geometry of D321G AEE with the non-native substrate.

RC

| | | | |
|---|---|---|---|
| C | 37.4909349 | 27.7310793 | 40.7492231 |
| H | 37.8359820 | 28.7259656 | 41.0601204 |
| H | 37.8240320 | 27.5417475 | 39.7223882 |
| N | 36.0148545 | 27.6838676 | 40.7882569 |
| H | 35.6495244 | 27.6262910 | 41.7800541 |
| H | 35.5865256 | 28.5316422 | 40.3598214 |
| H | 35.6131728 | 26.8893426 | 40.2413493 |
| C | 37.1149573 | 29.0102590 | 44.7727352 |
| H | 37.3026115 | 30.0719013 | 44.9840986 |
| H | 38.0597101 | 28.5707155 | 44.4106491 |
| C | 36.0571141 | 28.8667944 | 43.6607326 |
| O | 35.4934333 | 27.7642948 | 43.4789196 |
| O | 35.8915503 | 29.8788488 | 42.8976544 |
| C | 32.8940260 | 31.5397431 | 45.4904044 |
| H | 32.5332826 | 30.6565445 | 44.9431122 |
| H | 32.0766497 | 32.0525634 | 46.0158399 |
| C | 33.6448386 | 32.5612404 | 44.6014867 |
| O | 34.3016496 | 32.0739553 | 43.6160131 |
| O | 33.6092900 | 33.7435180 | 44.9357796 |
| C | 31.0816374 | 32.7194175 | 42.7841080 |
| H | 31.5822949 | 33.3775342 | 43.5112694 |
| H | 31.1750456 | 31.6865290 | 43.1491356 |
| C | 31.8153344 | 32.8678353 | 41.4404172 |
| O | 31.5174733 | 33.7932561 | 40.6708507 |
| O | 32.7364466 | 32.0014409 | 41.2006516 |
| C | 33.2572083 | 37.5149752 | 37.7522930 |
| H | 33.9379080 | 38.3484042 | 37.9854733 |
| H | 32.6774126 | 37.3054468 | 38.6666885 |
| C | 34.0410430 | 36.2807600 | 37.3209049 |
| H | 33.3635775 | 35.5044576 | 36.9242075 |
| H | 34.7385586 | 36.5240577 | 36.4949892 |
| C | 34.8250601 | 35.6627148 | 38.4690706 |
| H | 35.4813624 | 36.4196709 | 38.9358094 |
| H | 34.1220910 | 35.3468757 | 39.2598501 |
| N | 35.6476925 | 34.5103897 | 38.0948677 |
| H | 36.3944584 | 34.7921518 | 37.4507883 |
| H | 35.0575034 | 33.8481568 | 37.5849459 |
| Mg | 34.3273945 | 30.9558320 | 41.9564826 |
| O | 35.7684493 | 31.7204354 | 40.4831144 |
| C | 35.5047600 | 30.9041472 | 39.5545117 |
| O | 34.7374547 | 29.9256867 | 39.7846254 |
| C | 36.0834761 | 31.1060450 | 38.1259604 |
| C | 37.4870728 | 31.7330374 | 38.1133728 |
| C | 38.5392018 | 30.7065130 | 38.4710971 |
| C | 38.3840175 | 29.4384335 | 38.0424349 |
| C | 37.1802282 | 29.0427279 | 37.3199403 |
| O | 37.7715471 | 32.1907524 | 36.7861171 |
| C | 36.0694705 | 29.8291477 | 37.3061094 |
| C | 34.8795619 | 29.5138976 | 36.4776962 |
| O | 33.9409205 | 30.3036047 | 36.4055879 |
| C | 34.7966045 | 28.2291074 | 35.6621251 |
| C | 34.1208673 | 27.0691443 | 36.4162045 |
| C | 35.0270480 | 26.4762362 | 37.5013425 |
| O | 36.1473641 | 26.0269214 | 37.1177423 |
| O | 34.6143062 | 26.4905973 | 38.6862585 |
| H | 37.1718533 | 28.0831609 | 36.8014201 |
| H | 37.4971531 | 32.5793053 | 38.8082174 |
| H | 35.3946178 | 31.8264273 | 37.6616832 |
| H | 35.7883192 | 27.8957302 | 35.3296775 |
| H | 34.1999124 | 28.4869597 | 34.7733441 |
| H | 33.9026734 | 26.2772168 | 35.6797977 |
| H | 33.1705972 | 27.3867239 | 36.8632799 |
| H | 38.6736068 | 32.6024345 | 36.8119398 |
| H | 39.1843000 | 28.7085501 | 38.1908063 |
| H | 39.4538120 | 31.0381883 | 38.9704432 |
| O | 32.3319588 | 27.8510830 | 44.4351809 |
| H | 31.3684313 | 27.7684269 | 44.4651469 |
| H | 32.5710552 | 27.9936673 | 45.3797835 |
| H | 37.9554304 | 26.9884284 | 41.3978992 |
| H | 36.8942348 | 28.5258235 | 45.7238771 |
| H | 33.6144987 | 31.1871271 | 46.2284073 |
| H | 30.0319470 | 33.0112917 | 42.8161475 |
| H | 32.5618166 | 37.8356957 | 36.9766410 |

TS1

| | | | |
|---|---|---|---|
| C | 37.5172857 | 27.9184643 | 40.7750193 |
| H | 37.8270593 | 28.8872387 | 41.1912975 |
| H | 37.8991733 | 27.8422979 | 39.7487222 |
| N | 36.0400312 | 27.8445998 | 40.7483434 |
| H | 35.6450172 | 27.7071666 | 41.7147868 |
| H | 35.6074852 | 28.7199884 | 40.3696268 |
| H | 35.6618739 | 27.0891008 | 40.1249567 |
| C | 37.0963628 | 29.0170530 | 44.7789002 |
| H | 37.3013610 | 30.0732268 | 45.0012771 |
| H | 38.0276502 | 28.5643087 | 44.3991738 |
| C | 36.0102902 | 28.8948615 | 43.6884845 |
| O | 35.4688689 | 27.7892179 | 43.4750288 |
| O | 35.7990575 | 29.9392155 | 42.9808560 |
| C | 32.9142429 | 31.6405486 | 45.5279918 |
| H | 32.4912703 | 30.7926152 | 44.9704644 |
| H | 32.1380645 | 32.1865176 | 46.0819030 |
| C | 33.6994578 | 32.6345611 | 44.6386422 |
| O | 34.1975867 | 32.1716812 | 43.5684742 |
| O | 33.8317941 | 33.7956434 | 45.0527856 |
| C | 31.0553717 | 32.7718812 | 42.7747428 |
| H | 31.5499456 | 33.4301256 | 43.5068993 |
| H | 31.1735537 | 31.7372690 | 43.1253653 |
| C | 31.7698427 | 32.9496439 | 41.4275392 |
| O | 31.4606954 | 33.8981999 | 40.6797913 |
| O | 32.6869518 | 32.1026850 | 41.1574223 |
| C | 33.1154176 | 37.1723726 | 37.7571130 |
| H | 33.8759329 | 37.8956534 | 38.0899604 |
| H | 32.4668552 | 36.9555771 | 38.6210410 |
| C | 33.7616796 | 35.8852515 | 37.1973724 |
| H | 33.0183129 | 35.3282797 | 36.6021056 |
| H | 34.5686480 | 36.1577157 | 36.4910372 |
| C | 34.3330811 | 34.8877059 | 38.2140630 |
| H | 35.0683712 | 35.3659661 | 38.8776892 |
| H | 33.5323470 | 34.4708976 | 38.8474072 |
| N | 35.0142446 | 33.7573365 | 37.5166927 |
| H | 35.9145510 | 34.0598723 | 37.1155648 |
| H | 34.4058269 | 33.4413758 | 36.7479248 |
| Mg | 34.2559146 | 30.9929855 | 41.9529661 |
| O | 35.6339791 | 31.9422084 | 40.6504137 |
| C | 35.4614663 | 31.1289529 | 39.6692145 |
| O | 34.6997189 | 30.1182925 | 39.8946516 |
| C | 36.1199255 | 31.3740247 | 38.3726836 |
| C | 37.5009763 | 32.0292117 | 38.4913040 |
| C | 38.5969727 | 31.0031142 | 38.6405995 |
| C | 38.5027627 | 29.8488397 | 37.9430669 |
| C | 37.3010436 | 29.5352029 | 37.2033244 |
| O | 37.7538915 | 32.7786858 | 37.2949220 |
| C | 36.1256031 | 30.2258561 | 37.4037638 |
| C | 34.9203545 | 29.8259989 | 36.6226272 |
| O | 33.9091257 | 30.5241859 | 36.5951546 |
| C | 34.8891248 | 28.5537713 | 35.7668327 |
| C | 34.2074435 | 27.3593380 | 36.4594495 |
| C | 35.1011232 | 26.6931413 | 37.5120345 |
| O | 36.2054371 | 26.2308790 | 37.1146694 |
| O | 34.6749991 | 26.6561731 | 38.6960031 |
| H | 37.3280903 | 28.6786947 | 36.5292011 |
| H | 37.4937656 | 32.6967348 | 39.3693583 |
| H | 35.3827744 | 32.6162338 | 38.0051480 |
| H | 35.8812786 | 28.2279863 | 35.4317463 |
| H | 34.3058677 | 28.8316986 | 34.8746293 |
| H | 33.9864486 | 26.6083674 | 35.6813847 |
| H | 33.2593692 | 27.6558489 | 36.9238179 |
| H | 38.6794554 | 33.1307198 | 37.3357120 |
| H | 39.3303224 | 29.1330242 | 37.9318386 |
| H | 39.4772600 | 31.2415331 | 39.2444827 |
| O | 32.2944276 | 27.8488088 | 44.4339660 |
| H | 31.3349810 | 27.7382515 | 44.4930206 |
| H | 32.5560836 | 27.9966571 | 45.3718472 |
| H | 37.9575163 | 27.1243613 | 41.3780653 |
| H | 36.8834102 | 28.5291701 | 45.7300514 |
| H | 33.6258429 | 31.2312154 | 46.2450266 |
| H | 30.0002441 | 33.0427295 | 42.8123463 |
| H | 32.5002360 | 37.6414230 | 36.9892519 |

IM

| | | | |
|---|---|---|---|
| C | 37.5181407 | 27.9201286 | 40.7604117 |
| H | 37.7735183 | 28.8850236 | 41.2208265 |
| H | 37.9257945 | 27.9032101 | 39.7400952 |
| N | 36.0471871 | 27.7769614 | 40.7078999 |
| H | 35.6512974 | 27.6108214 | 41.6656797 |
| H | 35.5504905 | 28.6261888 | 40.3331228 |
| H | 35.7030932 | 27.0054835 | 40.0837548 |
| C | 37.0579883 | 29.0072233 | 44.7585377 |
| H | 37.2463214 | 30.0691885 | 44.9664601 |
| H | 37.9929894 | 28.5667067 | 44.3739667 |
| C | 35.9607290 | 28.8459613 | 43.6768745 |
| O | 35.4672965 | 27.7199588 | 43.4660806 |
| O | 35.6905457 | 29.8857871 | 42.9803943 |
| C | 32.9490614 | 31.6186645 | 45.4808410 |
| H | 32.5276267 | 30.7613154 | 44.9366854 |
| H | 32.1663098 | 32.1728197 | 46.0183600 |
| C | 33.7280876 | 32.6101348 | 44.5742323 |
| O | 34.1427657 | 32.1768382 | 43.4600637 |
| O | 33.9100492 | 33.7562063 | 45.0185397 |
| C | 31.0396605 | 32.7852440 | 42.7707217 |
| H | 31.5339049 | 33.4631954 | 43.4845090 |
| H | 31.1665976 | 31.7592833 | 43.1414680 |
| C | 31.7352086 | 32.9328472 | 41.4116698 |
| O | 31.4663609 | 33.9011144 | 40.6712660 |
| O | 32.5915142 | 32.0328043 | 41.1211630 |
| C | 33.1634601 | 37.3094921 | 37.8137595 |
| H | 33.9207058 | 38.0606422 | 38.0846648 |
| H | 32.5143744 | 37.1697925 | 38.6927644 |
| C | 33.8144503 | 35.9721302 | 37.4136387 |
| H | 33.0941519 | 35.3385837 | 36.8668931 |
| H | 34.6597119 | 36.1413589 | 36.7222842 |

C 34.2869346 35.1819695 38.6287639
H 35.0605199 35.7147685 39.2008947
H 33.4401510 34.9856389 39.3056171
N 34.8407763 33.8364667 38.2625152
H 35.8198124 33.7959223 37.8768201
H 34.2251990 33.3721012 37.5702053
Mg 34.1542771 30.9005599 41.8877134
O 35.4592431 31.8332701 40.4634319
C 35.3677089 30.8603101 39.5868074
O 34.5502130 29.9127175 39.9242999
C 36.2173972 30.8361718 38.4349449
C 37.4566732 31.6874686 38.4874657
C 38.6674724 30.8485108 38.1877285
C 38.5809199 29.7994660 37.3275662
C 37.3164264 29.3844725 36.7986987
O 37.3712735 32.8153729 37.5542903
C 36.1332012 29.8657227 37.3574601
C 34.8491571 29.5052085 36.6784006
O 33.8540408 30.2221448 36.7609600
C 34.7312294 28.2806746 35.7688243
C 34.0511803 27.0919172 36.4695021
C 34.9832616 26.4713934 37.5138301
O 36.0520227 25.9612420 37.0842606
O 34.6332399 26.5385621 38.7214954
H 37.2984776 28.6387195 36.0046931
H 37.5582571 32.1124821 39.4961369
H 34.8750077 33.2063629 39.0896506
H 35.6969751 27.9275321 35.3897355
H 34.1176895 28.6048275 34.9128077
H 33.8262495 26.3327867 35.7016803
H 33.1100960 27.3945183 36.9442373
H 38.2586436 33.2566786 37.5449378
H 39.4778401 29.2366644 37.0511701
H 39.6219606 31.1324220 38.6423305
O 32.2591753 27.7929602 44.4743795
H 31.3055111 27.6455320 44.5472851
H 32.5282725 27.9597260 45.4075103
H 37.9687075 27.1231117 41.3518682
H 36.8684765 28.5276387 45.7188247
H 33.6451352 31.2213544 46.2195514
H 29.9818856 33.0454385 42.8091675
H 32.5446965 37.7148200 37.0131922

TS2

C 37.5187003 27.8891706 40.7541037
H 37.7834564 28.8602330 41.1962219
H 37.9130559 27.8513013 39.7289198
N 36.0468712 27.7458528 40.7235508
H 35.6615285 27.5907172 41.6901881
H 35.5469121 28.5839874 40.3466177
H 35.6935002 26.9693804 40.1088075
C 37.0621166 29.0006881 44.7596509
H 37.2442542 30.0643402 44.9640345
H 38.0006783 28.5622186 44.3813892
C 35.9708845 28.8291742 43.6750785
O 35.4891181 27.6979505 43.4610201
O 35.6950470 29.8651890 42.9760666
C 32.9319441 31.6208399 45.4998057
H 32.5001293 30.7676168 44.9574443
H 32.1589231 32.1798902 46.0456686
C 33.7149416 32.6038132 44.5904175
O 34.1561122 32.1500145 43.4937131
O 33.8836987 33.7571858 45.0177227
C 31.0437196 32.7707029 42.7849016
H 31.5348667 33.4456126 43.5037938
H 31.1626447 31.7428874 43.1537657
C 31.7479973 32.9246861 41.4306418
O 31.4844186 33.8940012 40.6959950
O 32.6075614 32.0214459 41.1427718
C 33.2543176 37.3492505 37.7444994
H 34.0052500 38.1214029 37.9720224
H 32.6598775 37.1938862 38.6594096
C 33.9126352 36.0289377 37.3018697
H 33.1544140 35.3599046 36.8562669
H 34.6609004 36.2128021 36.5085530
C 34.5578907 35.2656698 38.4565091
H 35.4132927 35.8233495 38.8723555
H 33.8128529 35.1605621 39.2635301
N 35.0224768 33.9041457 38.0628381
H 36.3148553 33.5157419 37.7303767
H 34.3784796 33.5300107 37.3477167
Mg 34.1412781 30.8871336 41.9224088
O 35.4819226 31.7377770 40.4757681
C 35.3171487 30.8057041 39.5933098
O 34.4929324 29.8723301 39.9039320
C 36.1679782 30.7549098 38.4083058
C 37.4030447 31.5268832 38.4912839
C 38.5961341 30.8138590 38.0278827
C 38.4744162 29.7931416 37.1284203
C 37.1954517 29.3596513 36.6834861
O 37.2858514 32.9112201 37.5076275
C 36.0375331 29.8155030 37.3176307
C 34.7244852 29.4683287 36.6756000
O 33.7596680 30.2183962 36.7701136
C 34.5616694 28.2381083 35.7844436
C 33.9384902 27.0463915 36.5299743
C 34.9273572 26.4583655 37.5398733
O 35.9948227 25.9752933 37.0772723
O 34.6160593 26.5248353 38.7586268
H 37.1365434 28.6174243 35.8891337
H 37.5352672 32.0268629 39.4511095
H 34.8927603 33.2761971 38.8631554

| | | | |
|---|---|---|---|
| H | 35.5074933 | 27.8950446 | 35.3496404 |
| H | 33.8952316 | 28.5524896 | 34.9659925 |
| H | 33.6964865 | 26.2754750 | 35.7799476 |
| H | 33.0125358 | 27.3368399 | 37.0416447 |
| H | 38.1194157 | 33.4280458 | 37.6687335 |
| H | 39.3651039 | 29.2792087 | 36.7585765 |
| H | 39.5727038 | 31.1207782 | 38.4109500 |
| O | 32.2519357 | 27.7764036 | 44.4936530 |
| H | 31.2996367 | 27.6195620 | 44.5656143 |
| H | 32.5186357 | 27.9482396 | 45.4268064 |
| H | 37.9743507 | 27.0997529 | 41.3518241 |
| H | 36.8701967 | 28.5243385 | 45.7210691 |
| H | 33.6356441 | 31.2177061 | 46.2280615 |
| H | 29.9870303 | 33.0361984 | 42.8163167 |
| H | 32.5817710 | 37.7379748 | 36.9798808 |

PC

| | | | |
|---|---|---|---|
| C | 37.5003110 | 27.6820510 | 40.7132397 |
| H | 37.8231173 | 28.6685494 | 41.0664892 |
| H | 37.8360379 | 27.5542712 | 39.6752448 |
| N | 36.0273915 | 27.6153940 | 40.7696985 |
| H | 35.6870353 | 27.5271284 | 41.7726184 |
| H | 35.5555668 | 28.4687302 | 40.3815035 |
| H | 35.6083294 | 26.8313979 | 40.2315808 |
| C | 37.0342867 | 29.0000108 | 44.7658154 |
| H | 37.2003456 | 30.0661736 | 44.9690010 |
| H | 37.9725897 | 28.5657366 | 44.3829781 |
| C | 35.9324721 | 28.7926553 | 43.7002990 |
| O | 35.5755385 | 27.6219978 | 43.4264032 |
| O | 35.5077489 | 29.8235042 | 43.0852508 |
| C | 32.9084943 | 31.6746235 | 45.5540712 |
| H | 32.4116943 | 30.8415893 | 45.0367838 |
| H | 32.1856264 | 32.2679164 | 46.1317250 |
| C | 33.6739156 | 32.6202490 | 44.5916057 |
| O | 33.9476189 | 32.1545034 | 43.4469064 |
| O | 33.9541120 | 33.7532463 | 45.0047522 |
| C | 31.0307817 | 32.7706079 | 42.7696297 |
| H | 31.5348660 | 33.4920006 | 43.4297047 |
| H | 31.1516724 | 31.7694089 | 43.2036535 |
| C | 31.6873664 | 32.8262404 | 41.3817285 |
| O | 31.4970513 | 33.8112057 | 40.6498956 |
| O | 32.4133495 | 31.8171211 | 41.0623012 |
| C | 33.1932589 | 37.2721773 | 37.6616421 |
| H | 33.9877511 | 38.0058468 | 37.8722219 |
| H | 32.6320895 | 37.1196198 | 38.5980609 |
| C | 33.7770853 | 35.9443704 | 37.1467876 |
| H | 32.9691402 | 35.2951623 | 36.7578235 |
| H | 34.4551675 | 36.1229557 | 36.2920726 |
| C | 34.5214150 | 35.1427151 | 38.2081254 |
| H | 35.4151775 | 35.6961135 | 38.5483482 |
| H | 33.8548719 | 35.0368933 | 39.0877223 |
| N | 34.9413965 | 33.8244183 | 37.6987262 |
| H | 36.3472279 | 33.0981784 | 36.6672118 |
| H | 34.1106183 | 33.3560521 | 37.3150822 |
| Mg | 33.9470614 | 30.8285634 | 41.9756816 |
| O | 35.4559880 | 31.8676620 | 40.6869272 |
| C | 35.5305190 | 30.8378729 | 39.9607573 |
| O | 34.7091244 | 29.8889736 | 40.1248279 |
| C | 36.7301433 | 30.6212381 | 39.0730252 |
| C | 37.9605585 | 30.9496663 | 39.6558252 |
| C | 39.1647627 | 30.5660108 | 39.0629607 |
| C | 39.1478538 | 29.8738174 | 37.8541067 |
| C | 37.9283465 | 29.5956657 | 37.2331961 |
| O | 37.0591524 | 32.5029715 | 36.3101965 |
| C | 36.7071919 | 29.9548634 | 37.8208580 |
| C | 35.4371165 | 29.8147051 | 37.0301104 |
| O | 34.5259242 | 30.6012789 | 37.2081328 |
| C | 35.2895674 | 28.7174007 | 35.9786576 |
| C | 34.3435000 | 27.5955125 | 36.4402072 |
| C | 34.9465330 | 26.7959324 | 37.6053006 |
| O | 36.0730638 | 26.2694983 | 37.3885045 |
| O | 34.3140743 | 26.7440518 | 38.6935766 |
| H | 37.9284500 | 29.1212936 | 36.2517601 |
| H | 37.9666692 | 31.4811509 | 40.6088964 |
| H | 35.2144242 | 33.2394157 | 38.4901865 |
| H | 36.2531271 | 28.2618303 | 35.7204981 |
| H | 34.8694568 | 29.1995608 | 35.0828909 |
| H | 34.1977137 | 26.9034794 | 35.5923704 |
| H | 33.3643504 | 28.0119165 | 36.7052028 |
| H | 37.8246866 | 32.7168305 | 36.8614659 |
| H | 40.0829883 | 29.5657379 | 37.3812528 |
| H | 40.1036370 | 30.8151492 | 39.5540786 |
| O | 32.1780413 | 27.6973039 | 44.5303397 |
| H | 31.2381458 | 27.4944137 | 44.6438196 |
| H | 32.4688505 | 27.9111136 | 45.4479957 |
| H | 37.9772233 | 26.9358640 | 41.3487134 |
| H | 36.8562853 | 28.5241544 | 45.7301511 |
| H | 33.6253995 | 31.2426813 | 46.2523118 |
| H | 29.9744187 | 33.0364624 | 42.8081747 |
| H | 32.5091795 | 37.7056190 | 36.9321007 |
